# Integration of rate and phase codes by hippocampal cell-assemblies supports flexible encoding of spatiotemporal context

**DOI:** 10.1101/2023.12.06.570348

**Authors:** Eleonora Russo, Nadine Becker, Aleks P. F. Domanski, Timothy Howe, Kipp Freud, Daniel Durstewitz, Matthew W. Jones

**Affiliations:** The BioRobotics Institute, Department of Excellence in Robotics and AI, Scuola Superiore Sant’Anna, 56127 Pisa, Italy; Dept. of Theoretical Neuroscience, Central Institute of Mental Health, Medical Faculty Mannheim, Heidelberg University, 68159 Mannheim, Germany; Department of Psychiatry and Psychotherapy, University Medical Center, Johannes Gutenberg University, 55131 Mainz, Germany; School of Physiology, Pharmacology & Neuroscience, Faculty of Life Sciences, University of Bristol, University Walk, Bristol BS8 1TD, UK; Nanion Technologies GmbH, Ganghoferstr. 70A, D-80339 Munich – Germany; School of Computer Science, Merchant Venturers Building, University of Bristol, Woodland Road, Bristol, BS8 1UB, UK

**Keywords:** cell assemblies, precession, temporal coding, rate coding, CA1, theta phase, theta sequences

## Abstract

Spatial information is encoded by location-dependent hippocampal place cell firing rates and sub-second, rhythmic entrainment of spike times. These ‘rate’ and ‘temporal’ codes have primarily been characterized in low-dimensional environments under limited cognitive demands; but how is coding configured in complex environments when individual place cells signal several locations, individual locations contribute to multiple routes and functional demands vary? Quantifying rat CA1 population dynamics during a decision-making task, we show that the phase of individual place cells’ spikes relative to the local theta rhythm shifts to differentiate activity in different place fields. Theta phase coding also disambiguates repeated visits to the same location during different routes, particularly preceding spatial decisions. Using unsupervised detection of cell assemblies alongside theoretical simulation, we show that integrating rate and phase coding mechanisms dynamically recruits units to different assemblies, generating spiking sequences that disambiguate episodes of experience and multiplexing spatial information with cognitive context.

## Introduction

Hippocampal coding multiplexes over broad temporal scales incorporating prior, current, and future contextual information [1], [2]. Among pyramidal cells of hippocampal CA1, transient firing rate increases lasting from hundreds to thousands of milliseconds encode the position of an animal within the environment (‘place cells’ [3], [4]), routes through paths with overlapping segments (‘splitter cells’ [5], [6], [7]), signal goal-locations [8], mark time intervals [9], respond to specific odors [10], sounds [11], objects [12] and, in humans, to other people’s identities [13]. The information required to form these multimodal representations [14], converges on the hippocampus from cortical and subcortical regions [15], building context-specific cognitive rate- maps [16], [17].

In conjunction with rate coding, hippocampal units also coordinate at much faster timescales, entrained to the dominant state-dependent oscillations of the local field potential (LFP): 5-10 Hz theta rhythms during active exploration and REM sleep, and sharp wave-ripples (SWR) during immobility and non-REM sleep. Theta phase precession [18], theta sequences [19], and SWR-associated replay [20] produce single- or multi-unit activity patterns with temporal precision on the order of tens of milliseconds. During phase precession, the spike times of a place cell with respect to the ongoing theta oscillation shift to earlier phases in the cycle as the animal moves through that unit’s spatial receptive field (‘place field’). At the population level, the temporally ordered, sequential activation of place cells within a theta cycle gives rise to characteristic theta sequences. Both processes provide a temporal code for spatial information: during phase precession, the position of the animal within a cell’s place field correlates with the theta phase of that cell’s spikes [18], [21], while theta sequences reflect past and imminent trajectories [22], [23]. In addition to spatial information, recent studies have uncovered theta sequences reflecting sequences of events [24], current goals [25], [26], and hypothetical future experiences [27], suggesting contributions of hippocampal temporal coding to planning and speculation. There is also growing evidence that spikes fired during different relative phases of local theta cycles may encode different aspects of past and future experiences [28], [29].

Despite their different timescales, the information content and processes governing hippocampal rate and temporal coding are not independent [30]. Firstly, the order in which units activate during theta cycles and replays typically reflects the sequences in which place fields are crossed by the animal during exploration (but see also [31]), or sequences in which sensory cues are encountered [24]. Mechanistically, interplay between fast somatic inhibition and slow dendritic depolarization as the animal crosses the respective neuron’s place field has been proposed as a possible mechanism linking firing rate with phase precession [32], [33], [34], [35] that may be tuned by local inhibitory interneurons [36]. Whatever their mechanisms, despite prevailing evidence for cross-temporal dependencies, the functional implications of interactions between rate and phase coding for hippocampal information processing remain equivocal.

Here, we investigate and quantify the extent to which flexible and transient activation of place cell assemblies affects both firing rate and phase/temporal coding modalities of individual place cells, enabling the discrimination of different visits to the same locations under varied cognitive demands. We therefore used a method for unsupervised detection of functional assemblies that is able to identify the composition of detected assemblies alongside their characteristic coordination timescales. In particular, we aimed to quantify how the information encoded by rate assemblies at 100 ms - 1 s timescales can affect the < 100 ms temporal coding of their constituent units in the CA1 region of rats performing a spatial memory and decision-making task on a complex maze. Under these conditions, we found that both theta phase and firing rate of place cells shift when the cell activates within different assemblies recruited according to task trial demands. Rate and temporal codes therefore interact, allowing CA1 populations to parse repeated visits to the same locations into different cognitive contexts.

## Results

Six adult male Long Evans rats were trained to perform a spatial working memory decision-making task on an end-to-end T-maze [37], [38]. During each trial, rats learned to run from one side of the maze to the opposite to collect 0.1 ml of sucrose solution at reward locations. Trials were subdivided into free choice and guided runs. During choice runs, rats started from one of the two reward locations marked with ‘G’ in **Fig. 1a** and were directed by a moveable door to turn right (from G1) or left (from G2) into the central arm of the maze. Having traversed the central arm, rats had to choose whether to turn right or left at the open T-junction to continue towards the reward locations in ‘C’. A correct choice required rats to leave the central arm by turning in the same direction as they entered it (i.e. correct runs were from G1-C1 or G2-C2). Reward was delivered only upon correct trials. Choice trials were followed by guided trials that led the rats back to the ‘G’ side of the maze via a pair of predetermined turns guided by motorized moveable doors. All guided runs ended with reward. Data presented here are from rats that had learned task rules over between 16-23 days of habituation and training, and were performing 40 trials per recording session at between 71-90% correct. A total of 322 units was recorded from the dorsal CA1 (**Supplementary** Fig. 1) during 24 recording sessions from 6 rats. Among these, we isolated putative place cells by selecting units with a mean, on-maze firing rate between 0.2 Hz and 4 Hz and with spatial information above 0.5 bits/s [39]. The following analyses focus exclusively on the 218 units identified as putative place cells (with a median of 7, a minimum of 2, and a maximum of 20 putative place cells per session).

**Figure 1.**
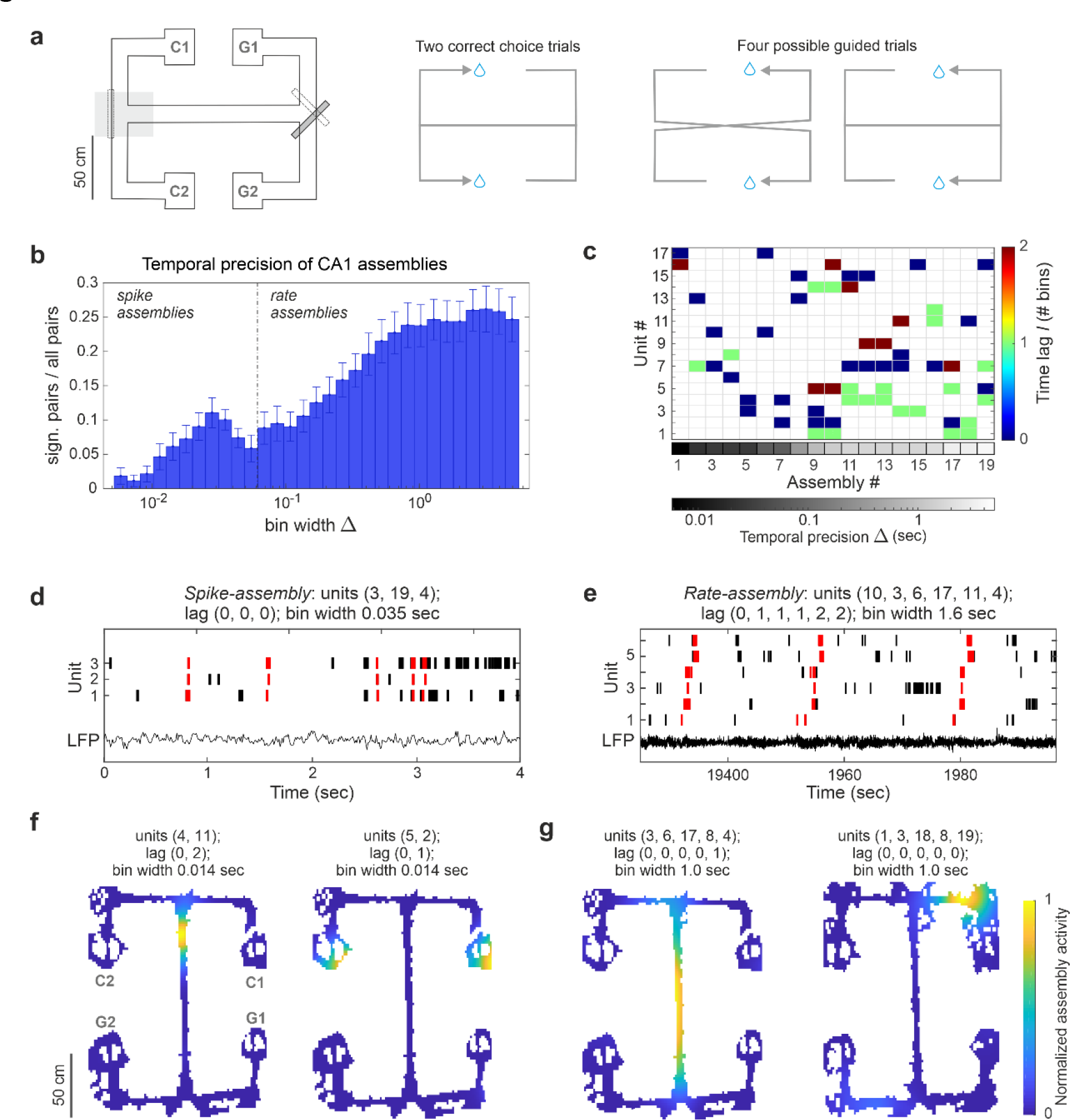
The two timescales of hippocampal assemblies. (a) Maze and task schematic. Task trials constitute choice and guided runs. In choice runs, the animal runs in direction G→C, deciding between left or right turns at the T-junction marked in gray. Correct choice is contingent upon starting location. In guided trials the animal runs in direction C→G, following a predetermined path guided by motorized moveable doors; (b) The distribution of the temporal precision of the assemblies detected during the spatial working memory task shows the presence of two predominant timescales: one peaked around 28 ms and one on the second scale. Bars show weighted mean and SE computed across animals and sessions. The mean is weighted by the number of place cells recorded in each session. See **Supplementary** Fig. 4 for the same analysis on assemblies detected excluding spikes fired during SWR; (c) Assembly-assignment matrix for one exemplary data set. The grayscale indicates the temporal resolution at which assemblies are detected; the color scale shows the lag between the activation of each assembly-unit with respect to the unit first active in the assembly. Units marked in dark blue (lag of 0) are the first to activate within the assembly, units marked in dark red the last (2 bins after the activation of the first assembly unit). Hippocampal place cell units were typically found taking part in multiple assemblies; (d, e) Examples of spike- (d) and rate- (e) assembly activity patterns (red) and raw LFP. Temporal resolution, composing units, and lag of activation of each unit with respect to the activation of the first assembly-unit are indicated in the figure. Spike-assembly activations appear to be locked to the theta rhythm of the LFP; (f, g) Example of spike- (f) and rate- (g) assembly activation maps (activity normalized to 1). See also **Supplementary** Fig. 3 for place fields of the relative assembly composing units.

Cell assemblies were identified with an unsupervised machine learning algorithm for Cell Assembly Detection (CAD) [40]. CAD detects and tests arbitrary multi-unit activity patterns that re-occur more frequently than chance in parallel single-unit recordings. The algorithm automatically corrects for non-stationarity in the units’ activities and scans spike count time series at multiple temporal resolutions, returning the characteristic timescales at which individual assembly patterns coordinate (**Supplementary** Fig. 2). Thanks to a flexible agglomeration algorithm, CAD can detect assemblies with any activity pattern, avoiding *a priori* suppositions about the characteristics of the detected motifs (see Methods). We use the term cell assemblies without making any assumptions about the anatomical connectivity between assembly units, which are identified solely based on their co-activation. Here we thus refer to a ‘functional cell assembly’ as any group of units whose activation coordinates with temporal precision between 5 msec and 5 sec, and arbitrary time lags between the unit activations, in a consistently reoccurring pattern.

### The two predominant timescales of hippocampal assemblies

As expected based on extensive previous analyses of place cell physiology, the temporal precision of hippocampal assemblies active during the task ranged from milliseconds to seconds and is bimodally distributed (Hartigan’s dip test for unimodality, n. bootstrap samples = 10^5^, dip=0.03, p = 0) into two major groups. We found: (1) 137 sharp spike patterns involving on average about 17% units per session per pattern (with a maximum of a 3-unit assembly in a 20-unit set) and a temporal precision in the range of 0.006 - 0.06 sec centered around 0.028 sec (*spike-assemblies*) and (2) 204 broader firing rate patterns with on average 28% units per session per pattern (with a maximum of an 11-unit assembly in a 19-unit set) and temporal precision between 0.07 - 5 sec (*rate-assemblies*) (**Fig. 1b**). This segregation into different timescales did not, however, correspond to different hippocampal cell populations; rather, spike- and rate-assemblies were composed of largely overlapping populations. About 83% of all assembly units participated in assemblies at both timescales. Moreover, two units taking part in the same spike-assembly were more likely to join the same rate-assembly than expected by chance (average probability of 0.9 against a chance level of 0.6, p < 10^-5^ computed by bootstrap, see Methods). Consistent with previous place cell analyses, this indicates that the same sets of hippocampal units coordinated at temporal precisions of both tens and hundreds of milliseconds [41], [42].

In order to understand the origin of these two characteristic timescales, we examined assembly activations in space and time. Assemblies are considered active whenever all units composing the assembly fire spikes matching the assembly activation pattern identified by the algorithm. The assembly is considered to have an activation of *n*, when all units composing the assembly fire at least *n* spikes in the bins matching the assembly pattern. **Figure 1** shows representative examples of activity patterns (**Figs. 1d, e**) and activation maps (**Figs. 1f, g**) for both assembly groups. Rate-assemblies reflected the simultaneous or sequential activation of the place fields of their constituent units in specific maze locations and/or along task-relevant trajectories, respectively (**Supplementary** Fig. 3). Their characteristic temporal scale, ranging from hundreds of milliseconds to seconds, was indeed compatible with the time needed by the animal to traverse the place field of a unit. Spike-assemblies, whose timescale is compatible with replay events or theta sequences [19], [20], had a more localized activation (with average spatial information of 2.67±0.09 in contrast to 1.91±0.07 for rate-assemblies, general linear mixed-effects model of the spatial information of assemblies according to the assembly type, spike- vs rate-assembly, F(1, 419) = 70.55, p = 7.0 10^-16^) which often appeared to be coordinated with the theta rhythm of the local field potential (**Fig. 1d**). To understand whether the observed coordination at short timescales was related specifically to replay events, we repeated the assembly detection but excluding epochs in correspondence of SWRs. As shown in **Supplementary** Fig. 4, the temporal resolution of the detected assemblies was conserved, showing that SWRs were not the prevailing source of fast coordination under these conditions.

### Spike-assemblies fire spikes phase-locked to the local theta rhythm

To quantify the relationships between assembly activations and the theta rhythm, we isolated the spikes fired by a unit while participating in assembly activations (*assembly-spikes*, in red in **Fig. 2a**) and compared their theta phase preference with the overall firing phase preference of that unit (in black in **Fig. 2a**). We found that both spike- and rate-assemblies were similarly phase modulated (**Fig. 2b**, generalized linear mixed-effects model of the probability of a unit phase-locking when firing within an assembly according to the assembly type, spike- vs rate-assembly; with binary dependent variable for significant phase locking and logit link function: F(1,1805) = 0.15, p = 0.70. The model accounts for rat identity and recording session as covariates. See **Supplementary** Fig. 5a for the same analysis excluding spikes fired during SWR), with about 70% of assembly-units phase-locked when active within an assembly configuration (all fractions presented here are computed after Benjamini–Hochberg correction for multiple comparisons, *α* = 0.05, see Methods for Hodges-Ajne test on phase locking). Yet, of these, the fraction of units with a stronger phase modulation during assembly activity when compared to their overall activity was significantly higher for spike-assemblies (75%) than for rate-assemblies (30%) (**Figs. 2c,d**, generalized linear mixed-effects model of the probability of a unit increasing phase modulation when firing within an assembly according to the assembly type, spike- vs rate-assembly; with binary dependent variable for significant increases in phase modulation and logit link function: F(1,1310) = 134.15, p = 1.3 10^-29^. The model accounts for rat identity and recording session as covariates. See **Supplementary** Fig. 5b for the same analysis excluding spikes fired during SWR). Thus, while theta-modulated units contribute to both spike- and rate-assemblies, rate-assembly activations did not specifically coincide with temporal windows of high theta modulation. On the other hand, the higher temporal precision of spike-assemblies was associated with enhanced phase-locking of their contributing members when active within the assembly configuration.

**Figure 2.**
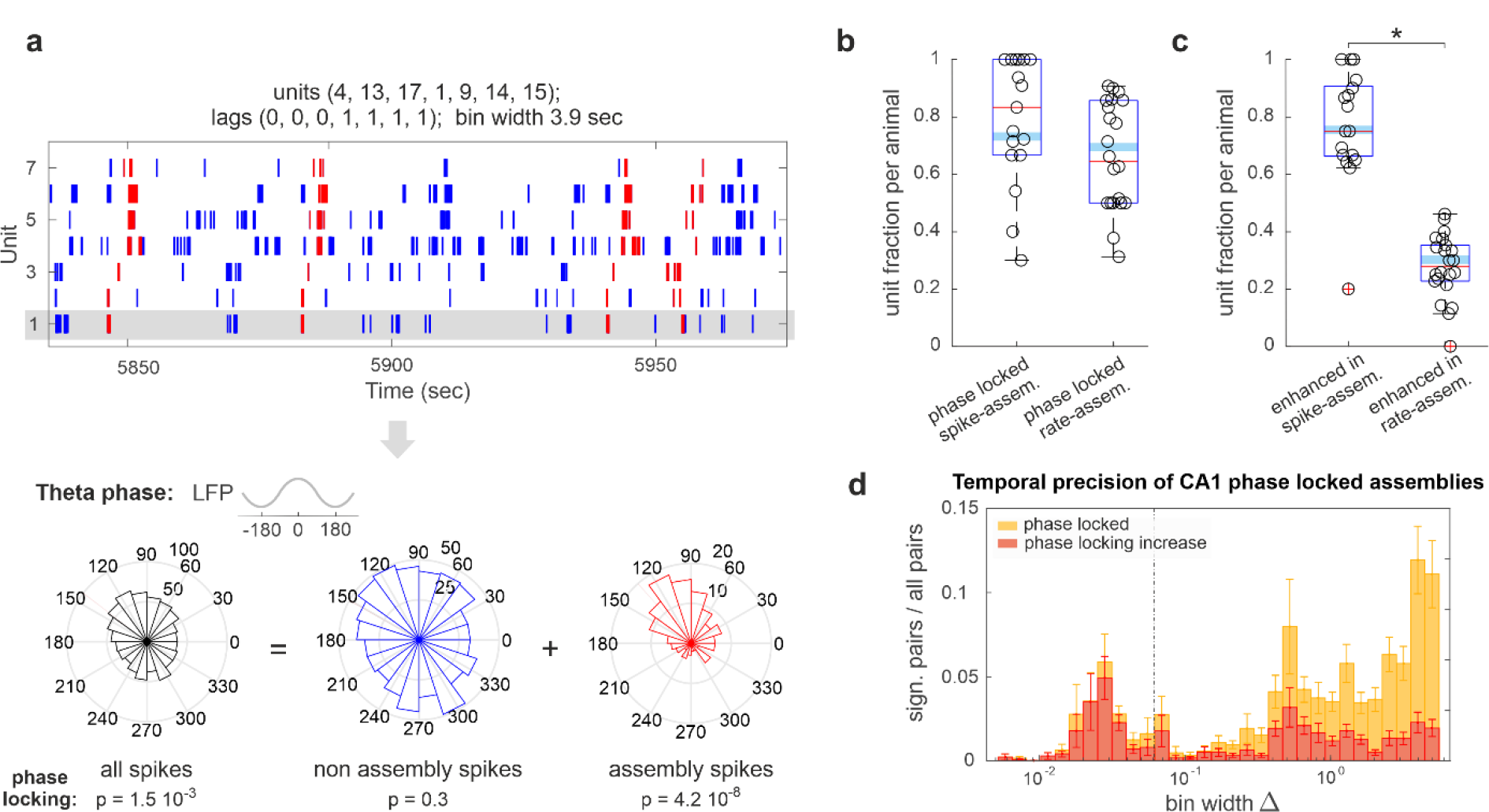
Spike-assemblies fire phase-locked spikes. (a) Raster plot of one typical assembly and its composing units (spikes fired during assembly activations marked in red). Below, phase histogram of the spikes of one exemplary unit (highlighted in gray) showing how assembly spikes (red) have an enhanced phase modulation with respect to either all (black) or non-assembly (blue) spikes; (b) Fraction of units with phase-modulated assembly-spikes and (c) units with assembly-spikes with enhanced phase-modulation with respect to the totality of the unit spikes (i.e. red vs. black in (a)) in at least one of the assemblies they take part in. Boxplots mark the median (red), the mean weighted by the number of tested assemblies per session (cyan), and the 25th and 75th percentiles (bottom and top edges of the box) computed across animals after Benjamini–Hochberg correction for multiple comparisons (*α* = 0.05), data points correspond to individual recording sessions; (d) Temporal precision of spike- and rate- assemblies with phase-modulated spikes (yellow) and assemblies with spikes with enhanced phase-modulation with respect to their composing units (red). Assembly pairs were detected with CAD*opti* separately in the 5 - 60 ms (spike-assembly) and 0.07 – 5.0 sec (rate-assembly) resolution window. CAD*opti* prunes redundant assemblies and selects those with the lowest p-value in each resolution window (see Methods). Bars show weighted mean and SE pooled from all sessions (mean weighted by the number of units per session). While the activity of both spike- and rate-assemblies is theta modulated, spike-assemblies in particular recruit spikes that have a stronger phase-locking than the totality of spikes fired by the unit.

### Individual units change phase preference when active in different assemblies

As single units were often contributors to multiple assemblies (cf. **Fig. 1c**) and assemblies activated with a characteristic phase preference (cf. **Fig. 2b**), we wondered whether the phase preference of hippocampal units is an assembly-specific property rather than a unit-specific one. In other words, do hippocampal units change their phase preference when active in different assemblies? We found that among all units taking part in at least two phase-locked assemblies (n = 64 in spike-assemblies and n = 169 in rate-assemblies) an average of 27% (fraction computed after Benjamini–Hochberg correction for multiple comparisons, *α* = 0.05) changed their phase preference when active in different assemblies of the same temporal resolution (**Figs. 3a, b,** see Methods for nonparametric test on equality of median phase). This relative *phase-shift* was found both in spike- and rate-assemblies, with a higher proportion of units with significant phase-shift in spike-assemblies (**Fig. 3b**, generalized linear mixed-effects model of the probability of a unit to phase-shift when firing in different assemblies according to the assembly type, spike- vs rate-assembly; with binary dependent variable for significant phase shift and logit link function: F(1,231) = 12.54, p = 4.8 10^-4^. The model accounts for rat identity and recording session as covariates. See **Supplementary** Fig. 5c for the same analysis excluding spikes fired during SWR).

**Figure 3.**
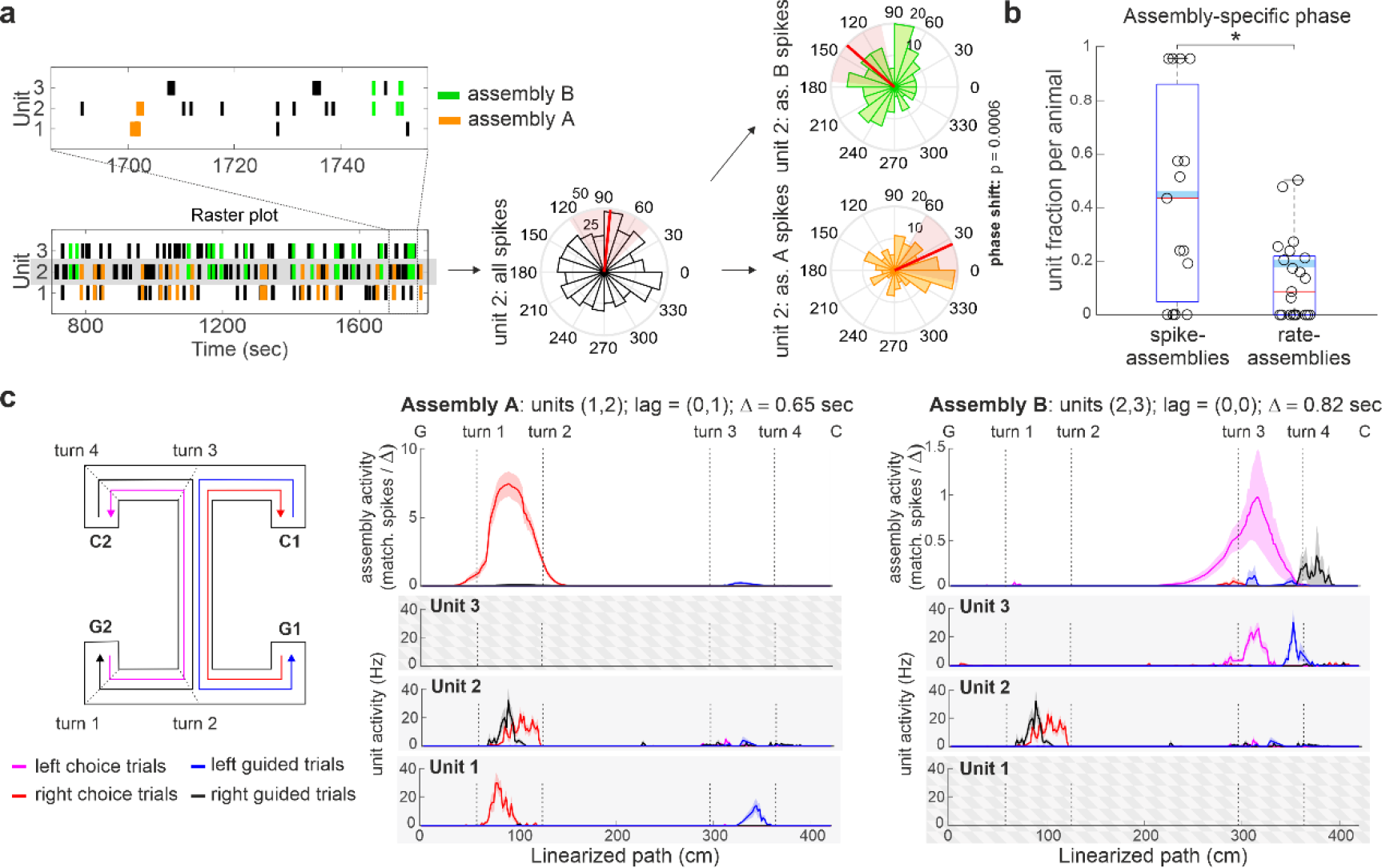
Single units change phase preference when active in different assemblies. (a) Raster plot and phase preference of two example assemblies, assembly A and assembly B, with a shared unit (unit 2). Unit 2 changes phase preference when active in the two different assembly configurations; (b) Fraction of units that change their phase preference when active in different assemblies. Boxplots mark the median (red), the mean weighted by the number of tested assemblies per session (cyan), and the 25th and 75th percentiles (bottom and top edges of the box) computed across animals after Benjamini– Hochberg multiple comparison correction (*α* = 0.05), data points correspond to distinct recording sessions; (c) Mean and SE activity along the maze of the two assemblies displayed in (a) (top) and their composing units during different trial types (bottom). The trial typologies displayed are: left (pink) and right (red) choice trials, and left (blue) and right (black) guided trials. The assembly and unit activity are plotted along the linearized path. Vertical dashed lines mark different task segments, along the path from C to G, as indicated in the maze scheme for the left choice trial. Assembly temporal resolution, composing units, and lag of activation of each unit with respect to the activation of the first assembly- unit are indicated in the figure title. Unit 2 takes part in both assembly A and B. When firing in the two assemblies, the unit fires preferentially at two different firing phases (a) to encode different task-related information (left vs. right choice trials). See also **Supplementary** Fig. 6 for other examples of assembly-dependent phase modulation of CA1 units.

### Phase coding of hippocampal assemblies

To investigate whether such shifts in phase encoded spatial or contextual information, we focused the analysis on those units changing phase when active in different contexts, comparing their activation during task epochs corresponding to different locations and cognitive demands on the end-to-end T-maze. **Fig. 3c** and **Supplementary** Fig. 6 show the activation along the maze of some typical units and the assemblies they joined. While single units fired in multiple locations, assembly activations were more selective (resulting in average spatial information of 2.25 ± 0.06 when compared with 1.52 ± 0.05 for single putative place cells), typically signaling one of the place fields of their constituent units and/or only a particular run type or direction.

This enhanced selectivity suggests that theta phase coding in hippocampal units extends beyond phase precession: while phase precession relative to the ongoing theta oscillation correlates with the distance covered by the animal within a unit’s place field, here we observed a phase preference disambiguating the activation of different assemblies in disjoined locations. We therefore hypothesized that theta phase coding of hippocampal units is not limited to the animal’s position within a place field, but is also associated with different locations or contexts in the environment.

### The theta firing phase of place cells can discriminate between distinct place fields of the same unit

A possible confound for the presence of contextual phase-shift coding in different assemblies comes from the fact that the spikes fired within an assembly might not uniformly sample the place field of a unit. Thus, if two assemblies systematically sampled the initial and final part of a phase-precessing unit’s place field respectively, this could result in an assembly-specific change in phase preference. To rule out this possibility, we analyzed the phase preference of single units, this time separating their spikes according to their own place fields instead of by assembly membership. In single units with multiple place fields, different rate-assemblies often activated in correspondence to different place fields of the unit (e.g. cf. Fig. 3c and Supplementary Fig. 6). As single units changed their phase preference when active in different rate-assemblies (cf. Fig. 3b), separating unit spikes by place field could reveal similar shifts in phase to those observed when separating them by rate-assembly.

Accordingly, we selected all units with multiple place fields and clustered their spikes according to their firing location (**Fig. 4a**, see Methods). We included in the analysis all recorded putative place cells, and not only those detected as participating in assemblies, as we assume that all CA1 place cells do participate in some assembly, but that sparse electrophysiological sampling of the CA1 population precludes the detection of all potential assemblies. In our dataset, putative place cells had more than one place field with a median of four fields per unit. Such a high number of place fields per unit was due to the relatively large size of the maze [43] and, critically, to its compartmentalization in clearly identifiable segments. Comparing the phase preference with respect to place field location, we found that 43% of the tested units changed firing phase when active in different place fields (value obtained as mean across sessions weighted by the number of units tested per session, 208 units among 24 sessions. Fraction computed after Benjamini–Hochberg correction for multiple comparisons, *α* = 0.05, **Figs. 4b, 5b**). Note that for this test we included all spikes fired within a place field and compared them with all spikes fired in another place field of the same unit. Thus, the observed phase-shift cannot be explained by a biased sampling of different subregions of the place fields.

**Figure 4.**
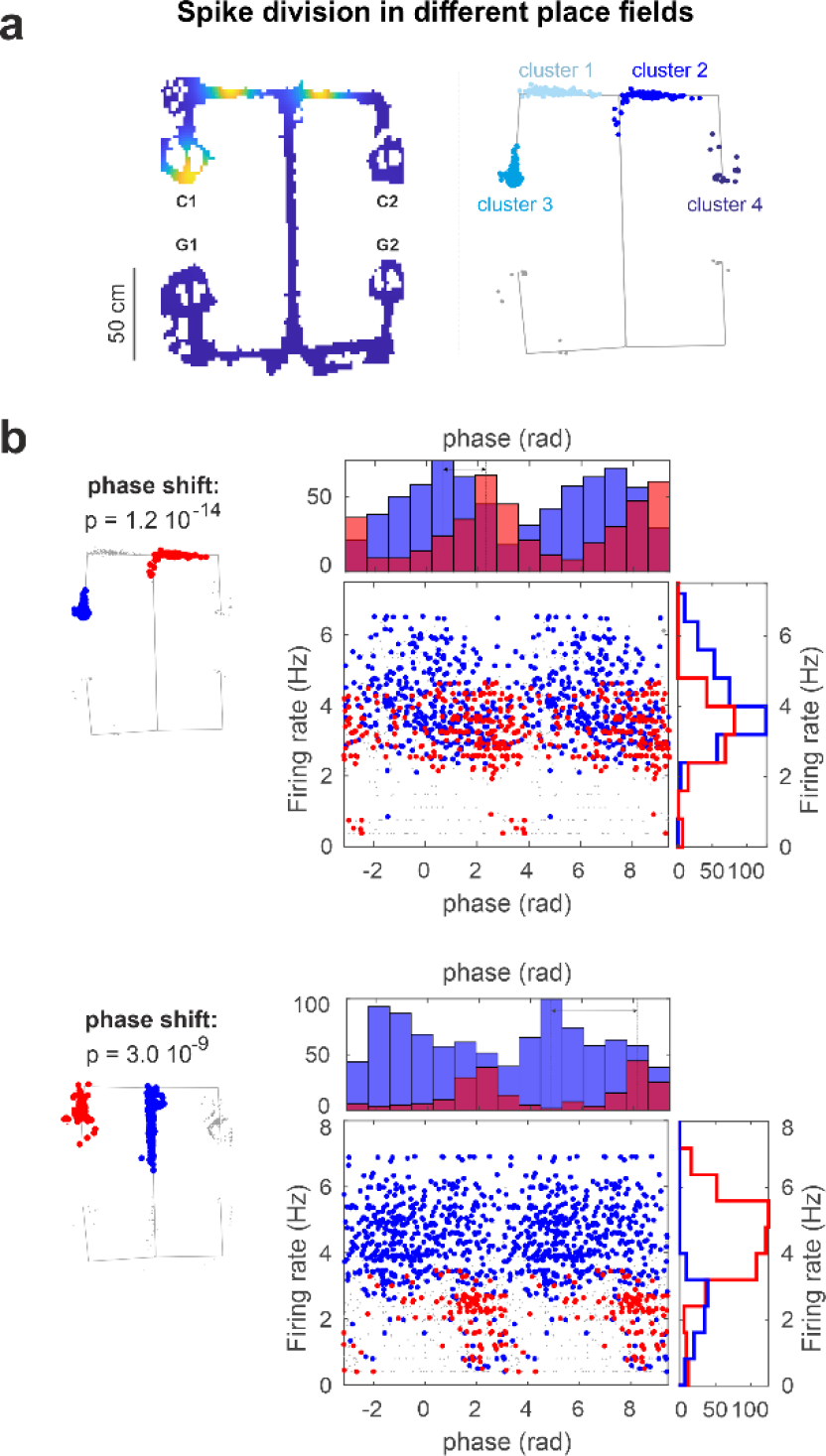
Place cells firing phase can encode distinct place fields. (a) Example of unsupervised spike clustering based on the spike position in a unit with multiple place fields. The number of place fields present in the spike set was established with DBSCAN, a density-based spatial clustering algorithm. The cluster memberships resulting from DBBSCAN were then fed to a Gaussian mixture model for the final step of the classification (see Supplementary Fig. 7 for a comparison between this and other methods for place field detection). (b) Rate-phase and respective marginal distributions of unit spikes color-coded according to their field of firing. The exemplary units changed their firing phase when active in different place fields. Grey dots are spikes fired out of the two tested place fields.

**Figure 5.**
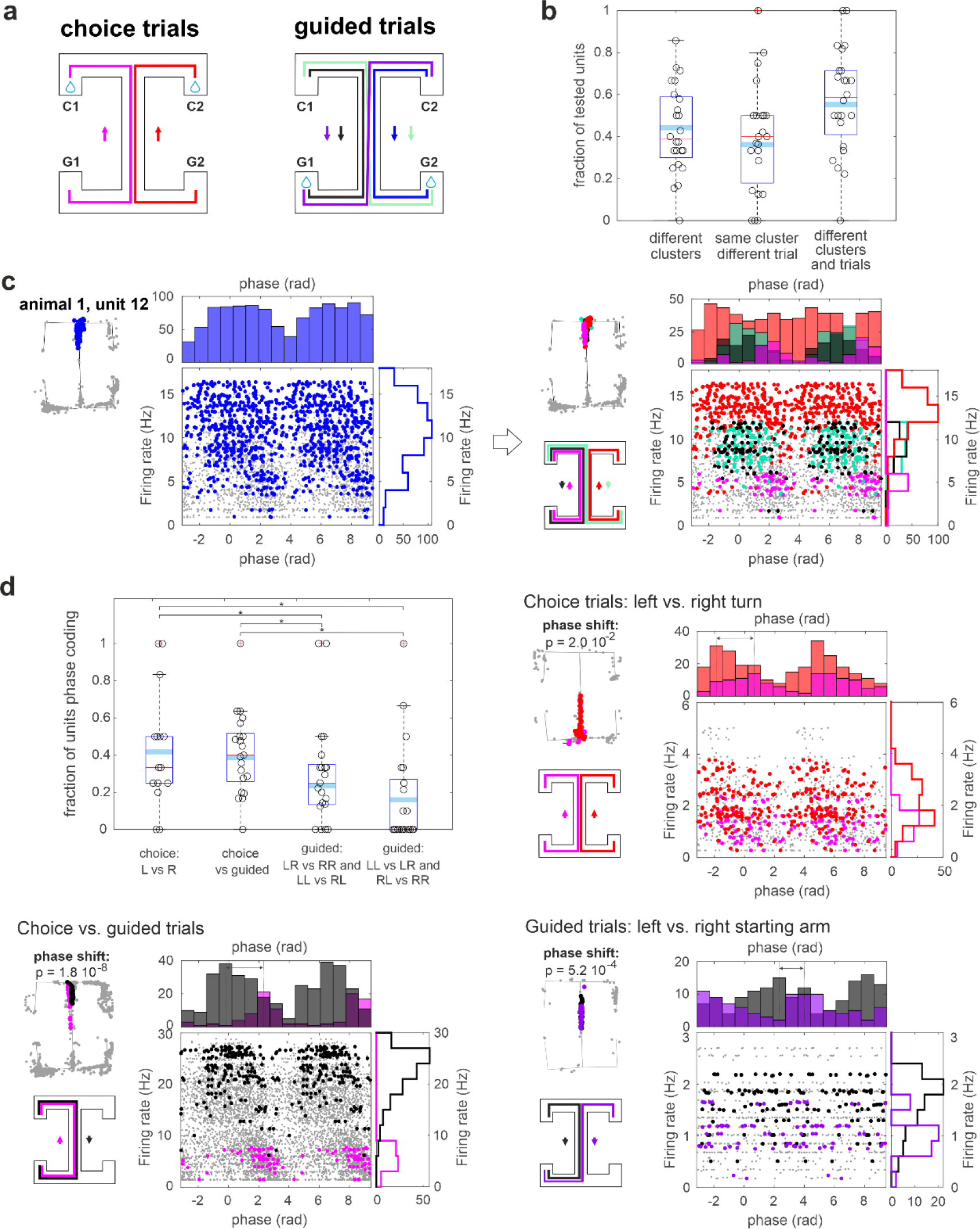
The firing phase of place cells can encode distinct task-related information within the same place field. (a) Trial categories: choice left (magenta), choice right (red), forced left (blue), forced right (black), forced switch right-left (light yellow) forced switch left-right (dark yellow); (b) fraction of units changing phase for different locations irrespective of the trial type, different trial types but same location, different trial type and/or location. See also Supplementary Fig. 8 and 9; (c) joint rate-phase distribution and marginal distributions for the spikes fired by a unit in one of its place fields (left panel, blue) and the same spikes divided by trial type (right panel, color coding of to trial type specified in (a)). Grey dots are spikes fired out of the two tested place fields; (d) fraction of units changing phase when comparing: left vs. right choice trials; choice vs. forced trials; forced trials with different origin arms; forced trials with same origin arms but different forced turn. See also Supplementary Fig. 10 and 11. Also displayed are example units changing phase preference (phase shift p-value indicated in figure) for the same place field during different trial types (same as (c)). In (b) and (d) boxplots mark the median (red), the mean weighted by the number of tested units per session (cyan), and the 25th and 75th percentiles (bottom and top edges of the box) computed across animals after Benjamini–Hochberg correction for multiple comparisons, (*α* = 0.05), data points correspond to distinct recording sessions.

### The firing phase of place cells can encode distinct task-related information within the same place field

Beyond encoding spatial information, the hippocampus has been shown to carry information about episodic memories [44], [45], sequences [46], [47] and abstract relations [48], [49]. Thus, hippocampal assemblies may encode task-relevant information beyond purely spatial parameters. We can therefore expect the recruitment of different assemblies when the animal has to remember, for example, a left turn rather than a right turn, or has to perform a guided turn rather than a choice turn. At the single-unit level, this should be reflected by a change in a unit’s phase preference for different trial types, even within the same location.

Separating unit spikes according to place field as in the previous analysis, we further divided the spikes according to the type of trial in which they occurred. Trials were divided into: left and right choice runs, when the animal had to choose between left and right turn; and four different types of guided runs, in which the sequence of turns was predetermined by the experimenter (**Fig. 5a**). We found that 36% of units changed their phase preference when active in different trial types above chance level, despite overlapping place-field locations (value obtained as mean across sessions weighted by the number of units tested per session, 160 units among 23 sessions. Fraction computed after Benjamini–Hochberg correction for multiple comparisons, *α* = 0.05, **Fig. 5b**, see also **Supplementary** Fig. 8 for same analysis with place fields identified with different methods and **Supplementary** Fig. 9 for same analysis with spikes fired only in epochs of high theta-power and no SWR). In fact, separating a unit’s spikes not just by place field but according to the task epoch during which they occurred resulted in narrower and more coherent phase distributions of a unit’s spikes (**Fig. 5c**). This observation was corroborated by training a support vector machine (SVM) classifier on the phase of spikes fired in an individual place field to distinguish between trial types. We found that for 32% of units at least two trial types could be distinguished above chance level within at least one of the unit place fields (see Methods for details). This result could not be explained by covariates such as the animal speed and the ongoing theta power (a general linear mixed-effects model of the accuracy of an SVM classifier trained on speed and theta power or on speed, theta power, and theta phase of each spike found a significant increase in accuracy for the latter classifier. Contrast tests between the two classifier types: F(1,530) = 8.9, p = 0.003. Test computed on the subset of place fields which could distinguish above chance trial identity in the latter, more powerful, classifier) nor the sorting cluster quality of the units (generalized linear mixed-effects model of a unit to phase-shift, either for different place fields or within the same place field but different trials, based on its L-ratio, recording session and animal; with binary dependent variable for significant phase change and logit link function: F(1,60) = 0.12, p = 0.74). Moreover, while it is known that splitter cells can differentiate between trial types by rate modulation, we found that adding information relative to the spike phase to the instantaneous firing rate further improved the decoding performance of the SVM (generalized linear mixed-effects model of the accuracy of an SVM classifier trained on the instantaneous firing rate of each spike or on instantaneous firing rate and phase of each spike. Contrast tests between the two classifier types: F(1,684) = 5.7, p = 0.017. Test computed on the subset of place fields which could distinguish above chance trial identity in the latter, more powerful, classifier).

Figure 5c shows an example unit with phase-shift coding at the choice junction of the maze, distinguishing left and right turns in choice trials and right turns in different guided trial types. Interestingly, in this example, the degree of differentiation of the unit phase is maximal when the animal has to actively remember its previous path to inform its next turn, and is absent in the guided trials when no active choice has to be made and the path covered from trial onset is identical (with a difference of 2.8 rad between the average spiking phase of the left and right choice trials, and of 0.2 rad between the two guided trials types). When examining instances of phase-shift coding among trial types, we found that the majority of phase-shifts distinguished between left vs. right choice trials and choice vs. guided trials (Fig. 5d, generalized linear mixed-effects model of the probability of a unit to change firing phase within the same place field according to the trial type, with binary dependent variable for significant phase change and logit link function: F(4,641) = 10.2, p = 4.9 10^-8^; contrast tests between specific condition/bars (I vs II) F(1,641) = 0.2, p = 0.6; (I vs III) F(1,641) = 5.6, p = 1.8 10^-2^; (I vs IV) F(1,641) = 10.7, p = 1.2 10^-3^; (II vs III) F(1,641) = 9.8, p = 1.8 10^-3^; (II vs IV) F(1,641) = 16.4, p = 5.8 10^-5^; (III vs IV) F(1,641) = 1.9, p = 0.2. The model accounts for rat identity and recording session as covariates). These results withstood using different methods to identify place fields (**Supplementary** Fig. 10) and removing spikes fired during epochs of low theta-power or during SWR (**Supplementary** Fig. 9). This further supports the link between phase-shift and information encoding. The type of trial was, in fact, the most relevant information for performing the task correctly.

### Phase-shift leads to context-dependent fine temporal coordination among units

The firing phase of hippocampal CA1 principal cells can therefore encode task related information to differentiate distinct maze locations or distinct mnemonic information within the same location. Such phase coding goes beyond what would be expected by phase precession. Nevertheless, phase-shifting and phase precession could share common underlying mechanisms. We observed that the population of units with phase-shift coding in at least one of their place fields correlated with the population of units with phase precession in at least one of their place fields (chi-square test for independence: *χ*^2^(1, *N* = 213) = 7.3, *p* = 6.8 10^−3^, p-value threshold for significance on the test for phase precession and phase-shift of 0.05, see Methods). Phase-shifting could occur between place fields with or without phase precession (Figs. 6a, b, c). When place-field-dependent phase-shift and phase precession co-occurred, the spike phases covered during the precession spanned different phase ranges in the two different fields (Fig. 6c).

**Figure 6.**
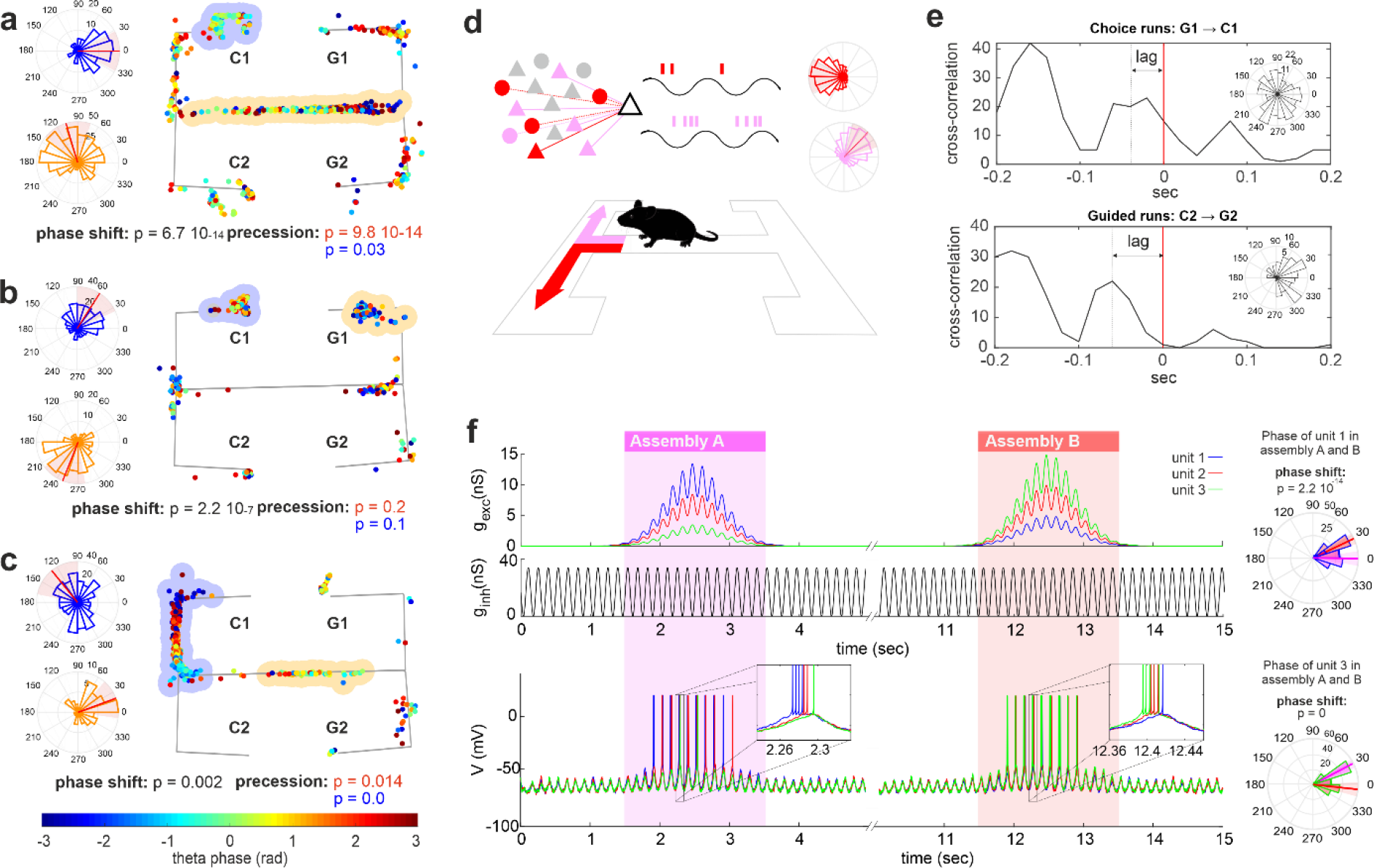
Phase-shift leads to context-dependent fine temporal coordination among units. (a) Spike phase (color-coded) and location of three units with multiple place fields. On the left, phase histogram of all spikes fired in the place fields marked in blue and yellow. All three units show a change in phase preference when active in different locations. Phase-shift could occur between place fields with (a-c) and without (b) phase precession. See Methods for tests on phase-shift and phase precession, p-values indicated in the figure; (d) Schematic of assembly-specific phase coding: In order to perform a task, or a choice, we are required to recall past information and future goals. Different contextual conditions will therefore trigger the retrieval of different assemblies even within the same location. By providing different degrees of depolarization, the retrieval of each assembly will set for their member-units assembly-specific ranges of firing phase; (e) Cross-correlation between the spikes of two units, unit *n* and unit *m*, during choice (above) and guided (below) right trials. Unit *n* changes firing phase when active in choice and guided trials (p = 0.03, phase histograms of *n*’s spikes are shown as inset for the two trial types) thereby changing its relative lag of activation with unit *m* during theta cycles (two-tailed Wilcoxon rank sum test, p = 0.01). Phase-shift coding can therefore produce context-dependent theta sequences. (f) Simulation of the recruitment of three neurons by two rate assemblies. (Top) Excitatory and inhibitory conductances of three neurons (blue, red, and green) modeled with an adaptive exponential integrate-and-fire model. The activation of rate assembly A and B was modeled with a transient unit-specific increase in excitatory conductances *g*_*exc*_ and a shared oscillatory inhibition *g*_inh_. (Bottom) Membrane potential *V* of the simulated units during assembly retrieval. (Right) Phase histogram of the spikes of units 1 (blue) and 3 (green) when active in assemblies A (magenta) and B (red). Spike’s phases are computed with respect to the oscillatory modulation of *g*_*exc*_. The change in assembly-specific depolarization provided to the simulated units leads to a change in the preferred phase of firing of the units (right) and, at the population level, in the order of unit activation along the theta cycle (inset).

Both experimental and theoretical studies have shown how a change in excitation received by a hippocampal unit can modify its phase of discharge [32], [33], [34], [35], [36], [50]. This suggests a possible interpretation of the phase-shift phenomena: spatial exploration or cognitive tasks recruit specific and diverse cell (rate-)assemblies; each assembly is characterized by a set of synaptic connections that provides the assembly-units with a characteristic level of excitation whenever the assembly is activated upon a specific task event; this assembly-specific degree of depolarization, combined with an oscillatory somatic inhibition, would thereby generate a phase preference, or phase-range preference, typical and specific for the activity of the unit within that assembly (Fig. 6d). In line with this hypothesis, we found that differences in average phase preference between two sets of spikes also co-occurred with differences in the average instantaneous firing rate (see Methods for methodological details). This was true when comparing spikes from different place-field locations, within the same location but from different trial types, or spikes occurring as part of different spike- or rate-assemblies (chi-square test of independence on pairs of spike-sets with p-value > or < 0.05 when testing phase and instantaneous firing rate differences: different place fields: *χ*^2^(1, *N* = 2165) = 28.9, *p* = 7.5 10^−8^, same location different trial types: *χ*^2^(1, *N* = 1004) = 20.5, *p* = 5.9 10^−6^, spike-assemblies: *χ*^2^(1, *N* = 318) = 4.9, *p* = 0.027, rate-assemblies: *χ*^2^(1, *N* = 5078) = 63.2, *p* = 2.0 10^−15^).

Finally, as for phase precession, the fine coordination of neuron firing with the theta oscillation will generate, at the network level, a stereotypical sequence of unit activations [19]. In the case of phase-shift coding however, the sequence of active units during a theta cycle will be determined not only by the spatial proximity of their place fields, but will also be affected by the identity of the assembly recruited at the time. This implies that the lag between the activation of two units within the theta cycle, and consequently the sequence order and composition, could vary according to the cognitive demand. We investigated this hypothesis by testing if the lag of maximal cross-correlation between two units within a theta cycle changed in different trial types (Fig. 6e). To make sure that such a change was not due to occasional outliers but was consistent across trials, we selected the lag of maximal cross-correlation for each trial and compared the set of lags so obtained by trial type. We performed the analysis for each place field of each unit and found that changes in cross-correlation lags occurred with higher probability in place fields where units displayed phase-shift coding across trials (chi-square test of independence performed on place fields according to DBSCAN + GMM, *χ*^2^(1, *N* = 11305) = 95.8, *p* = 0; DBSCAN, *χ*^2^(1, *N* = 11305) = 63.6, *p* = 1.6 10^−15^; RM + GMM, *χ*^2^(1, *N* = 8844) = 172.7, *p* = 0; RM, *χ*^2^(1, *N* = 9747) = 113.1, *p* = 0; also confirmed when excluding spikes fired during SWR and low theta-power epochs: DBSCAN + GMM, *χ*^2^(1, *N* = 11305) = 62.9, *p* = 2.2 10^−15^; DBSCAN, *χ*^2^(1, *N* = 11305) = 69.2, *p* = 1.1 10^−16^; RM + GMM, *χ*^2^(1, *N* = 8844) = 84.0, *p* = 0; RM, *χ*^2^(1, *N* = 9747) = 25.3, *p* = 4.8 10^−7^).

To explore how the activation of different rate assemblies can produce different activation sequences along the theta cycle even within the same set of units, we simulated the phenomenon with an adaptive exponential integrate-and-fire model ([51]; see Methods for formal description of the model)). As observed in the experimental data, single units could participate in multiple assemblies (cf. Fig. 3) and respond at different, but context-consistent, average firing rates (cf. Fig. 5). We simulated three units taking part in two rate assemblies. During the assembly activation, each unit received an assembly-specific degree of depolarization. Assembly retrieval was modeled by the synchronous and transient depolarization of its constituent units, while inhibition was on average constant over time, and identical for all units and both assemblies. Similarly to soma-dendritic interference models [32], [50], we captured the interplay between the oscillatory inputs to units by sinusoidally modulating both excitatory and inhibitory conductances with a relative offset of π rad (Fig. 6f **top**). As expected, assembly activations produced a broad increase in unit firing rate punctuated by faster temporal coordination with the theta oscillation (Fig. 6f **bottom**). This coordination was assembly-dependent and the simulated units changed their phase preference of firing, here computed with respect to the timecourse of the excitatory depolarization, when active in the two assemblies (Fig. 6f **right**). To formally detect multi-unit activity patterns generated by the assembly activations, we simulated multiple retrievals of the two assemblies and analyzed the obtained spike trains with CAD*opti* assembly detection algorithm, as previously done with our experimental dataset (see Methods and **Supplementary Table 1** for simulation details and CAD*opti* parameters). The activation of both assemblies produced, at a fine temporal scale, sequential activity patterns with a mean lag of 0.06 ± 0.005 sec and 0.07 ± 0.005 sec of the second and the third unit from the first active. Despite the similarity in pattern structure, the two assemblies triggered a different activation order of their constituent units. In 98% of the simulations (n = 100 simulations performed with different noise realizations), the detected activation sequence reflected the assembly-specific degree of depolarization provided to each unit (Fig. 6f **inset**). More generally, units that received the highest depolarization activated first within the theta cycle while, importantly, units receiving just a light depolarization terminated the activation sequence.

Overall, these results highlight how systematic shifts in theta phase preference not only broaden the range of information encoded by the hippocampal temporal coding but also contribute to the generation of context-specific theta sequences.

## Discussion

This study extends analyses of phase coding by CA1 place cell assemblies to a spatial working memory and decision-making task on a complex maze. We detected place cell assemblies using an unsupervised algorithm able to extract coordinated activity from the data without pre-defining timescales of interest. This corroborated extensive evidence that hippocampal coding is characterized by two predominant time scales: a rate scale (‘rate- assemblies’) reflecting place field firing rate modulation, and a sub-second temporal scale (‘spike-assemblies’) compatible with the entrainment of spikes by theta rhythms and during SWR-associated replay events. The relatively broad timescale characteristic of rate- assemblies suggests that their coordination is not solely imposed by the shared modulation of their composing units by the local theta rhythm, in agreement with their robustness to degradation of cholinergic signaling [52]. Nevertheless, spikes fired within either spike- and rate-assemblies coordinated with the theta rhythm, in line with previous literature reporting theta phase-locking of CA1 neurons when active in an assembly configuration [53], [54], [55]. We show that such theta locking is most pronounced for spike-assemblies, but present in rate- assemblies as well. A possible explanation for this enhanced phase locking is that isolating spikes fired within an assembly configuration effectively separates a specific mode from the otherwise multimodal phase distribution of the unit firing. In fact, the enhanced phase modulation of hippocampal units during assembly activations also revealed that units taking part in multiple assemblies changed their preferred spiking phase according to the assembly active at the time. This shift often coincided with a change in the location of place-field of activation of the unit when active in one or the other assembly. The observed higher specificity of information coding by assemblies with respect to that of the participating units therefore agreed with the higher specificity in phase preference exhibited by single units when active as part of the assembly. This was true both for rate- and, more frequently, spike-assemblies, and was not induced by SWR-associated replay events.

The enhanced phase modulation of hippocampal units during assembly activations also revealed that units taking part in multiple assemblies changed their preferred spiking phase according to the assembly active at the time. This modulation, not attributable to replay events, was validated at the single-unit level by grouping spikes by the place fields in which they occurred, rather than by assemblies. In the dorsal CA1’s deep sublayers, place cells can exhibit dual theta-phase firing preferences as the animal crosses the cell’s place field [28], [56], [57]. Here we show that changes in preferred firing phase can distinguish between distinct place fields or between different visits to overlapping locations on alternative routes, particularly under conditions that require active use of spatial memory and/or decision-making. These changes were fast and reversible, in line with the hypothesis that they were generated by the transient activation of different cell assemblies.

Our findings may reflect mechanisms related to those driving theta phase precession in CA1 place cells. Although unanimous agreement about those mechanisms is yet to be reached, models fall into three broad categories: interference between oscillations of the somatic membrane potential and of dendritic potentials at a slightly higher frequency [18], [58]; progressive dendritic depolarization coupled with somatic oscillatory inhibition discharge [32], [33], [34], [35], [50]; and patterns of synaptic transmission delays [59]. To test these various hypotheses, numerous experimental efforts have been made to elucidate the relationship between cell depolarization and spiking phase. Spatially uniform inhibitory conductance has been shown to enhance the range of phase precession [36]. It has been observed that while the animal moves towards the center of a cell’s place field, rate and phase of spiking strongly correlate [32], [35]. This correlation is, however, lost as the rate peak is passed and the animal leaves the cell’s place field [60], [61]. In vivo whole-cell recordings have shown that, during phase precession, the baseline membrane potential of CA1 pyramidal neurons undergoes a ramp-like depolarization [62]. In vitro, whole-cell patch-clamp recordings from dendrites and somata showed that an increase in dendritic excitation, coupled with phasic somatic inhibition, causes an increase in the neuron’s firing and the advancement of the spiking phase with respect to the somatic modulation [34], [50]. Similar results were observed when progressively depolarizing the membrane potential of hippocampal cells in anesthetized animals [33].

In line with this evidence that changes in depolarization lead to changes in discharge phase, we found in our data that the changes in phase preference of individual units also co-occurred with changes in the instantaneous firing rate. While a recent study has shown that anatomically distinct place cell subpopulations in superficial and deep sublayers of dorsal CA1 pyramidal cell layer bias towards rate and phase coding of spatial information respectively, as the richness of sensory cues in the local environment is experimentally manipulated [63], our results suggest an integration of rate and phase coding *within* a population to supports coding of complex information. This was also replicated through a model of adaptive exponential integrate-and-fire units similar in spirit to the soma-dendritic interference models ([32], [50]; c.f. Fig. 1e). The model confirmed that the activation of an assembly imposed an assembly-specific rate and phase preference on each assembly unit, thus producing, at the population level, assembly-specific theta sequences. Thus, while correlation between rate and phase changes has been observed during rate remapping [64], our findings demonstrate that phase-shift coding extends beyond rate remapping and occurs also between distinct place fields or assemblies, frequently coinciding with changes in instantaneous firing rate, similar to those observed for splitter cells [5], [6], [7]. The model predictions are also in line with recent work showing that individual place cells quickly switch between the encoding of alternative future locations or heading directions on alternate theta cycles, with lower firing rates for non-preferred directions occurring in later theta phases [27]. The mechanism highlighted by our analysis and model is not the sole modulator of firing of CA1 cells, as the timing of specific inputs from Schaffer collaterals and the perforant path also plays a role in their phase of discharge [56], [65]. Nevertheless, it captures the role played by various depolarization settings in inducing a modulation and discretization of phase preference of hippocampal CA1 place cells. We therefore propose that phase-shift coding may be a consequence of the different levels of depolarization generated by the specific constellations of synaptic input resulting from the activation of different assemblies as the animal encounters different cognitive or environmental contexts (cf. Fig. 6d).

The fine temporal coordination of unit activities imposed by phase precession is commensurate with the induction of plasticity mechanisms for binding episodic information that, otherwise, would be separated by seconds [66]. For this reason, phase precession has been proposed as a network mechanism for episodic sequence learning [1], [19], [35], [67], [68], [69], [70]. In support of this hypothesis, degradation of the temporal coordination of hippocampal units with the theta rhythm, e.g. by the administration of the cannabinoid receptor agonist [71] or by muscimol injection into the medial septum [72], led to reduced performance in memory tasks despite leaving place-field representations intact. Degraded phase precession caused by the passive transportation of rats during spatial exploration also drastically reduced replay during subsequent sleep [73], potentially reflecting impaired memory consolidation [74], [75], [76]. The changes in assembly-specific phase preference of hippocampal units reported here allow rapid reconfiguration of different theta sequences within the same neural population, differentiating distinct maze locations or distinct mnemonic information. This could contribute to the formation of context-dependent theta sequences [77], supporting the formation of episodic memories and planning [27].

Goal-dependent theta sequences have been observed during decision-making tasks, where the theta sequences terminated with the activation of cells encoding distant goal locations [26]. The process by which goal-related theta sequences form is still unclear, as is the causal relation between phase precession and theta sequences (see [78] for review). One hypothesis is that during the early stages of learning, inter-regional assemblies (e.g. prefrontal-hippocampal assemblies [77], [79], [80], [81], [82], [83] or medial-septum-hippocampal assemblies [84]), recruited at each theta cycle [85], modulate the depolarization of hippocampal units thereby dynamically producing goal-dependent cycling activity. In particular, as suggested by our theoretical model (cf. Fig. 6f), a low-level generalized depolarization of the cells encoding the current goal location could explain their spiking at the end of theta sequences, even when the animal is far from that location [26]. Similarly, the cognitive segmentation of a task-induced by the presence of landmarks and corners within a maze, could induce a shared enhanced depolarization to all the cells involved in the same ‘cognitive segment’, also in those with place field far from the animal but within the traveled maze segment. This could give rise to the observed space chucking of hippocampal theta sequences [22], [26]. Finally, similar mechanisms as observed here for assembly-dependent phase preference of CA1 units may also play roles in organizing phase preferences of neurons across other brain regions, consistent with evidence for phase precession in the dentate gyrus [86], CA3 [86], [87], entorhinal cortex [86], [88], subiculum [89], ventral striatum [90] and in the medial prefrontal cortex [38]. Such distributed processing would therefore support the integration of spatial and temporal information into cognitive contexts at a timescale commensurate with rapid adaptive behaviors, dynamically aligning different hippocampal assemblies with different subsets of neocortical and subcortical neurons.

## Acknowledgments

We would like to thank Richard J. Gardner for spike sorting, Andreas Draguhn and Martin Both for their support and the stimulating discussions, and Thomas McHugh and David Foster for commenting on the manuscript. The drawing of the rat in Figure 6 has been modified from https://scidraw.io/. ER has been supported by #NEXTGENERATIONEU (NGEU) and funded by the Ministry of University and Research (MUR), National Recovery and Resilience Plan (NRRP), project MNESYS (PE0000006) – A Multiscale integrated approach to the study of the nervous system in health and disease (DN. 1553 11.10.2022), by the Boehringer Ingelheim Foundation grant “Complex Systems”, and by the Ch. and H. Schaller Foundation. DD was supported by the Deutsche Forschungsgemeinschaft (DFG) within CRC-1134 (subproject D01) and through individual grant Du 354/10-1. Data acquisition was supported by BBSRC grant BB/G006687/1 awarded to MWJ and a Newton International Fellowship awarded to NB, with further analyses supported by a Wellcome Senior Research Fellowship to MWJ (202810/Z/16/Z).

## Author contributions

Experimental design and Electrophysiology, NB, TH and MWJ; Analysis, Visualization and Interpretation, ER; Writing - Original Draft, ER; Writing - Review and Editing, ER, MWJ, APFD, KF and DD; Resources, MWJ, ER and DD; Funding Acquisition, ER, NB, MWJ, and DD.

## Data availability

All data reported in this study are available from the corresponding authors upon request.

## Code availability

Code for cell assembly detection at multiple timescales CAD and CAD*opti* available at https://github.com/DurstewitzLab/Cell-Assembly-Detection and https://github.com/DurstewitzLab/CADopti, respectively.

## Methods

### Animals and husbandry

All procedures were conducted in accordance with the UK Animals (Scientific Procedures) Act 1986 and approved by the University of Bristol Animal Welfare and Ethical Review Board. Six adult male Long Evans rats (300–500g, Harlan, UK) were used in this study. Prior to surgery, rats were group-housed on a 12/12 hour light/dark cycle (lights on from 07:00–19:00) with free access to food and water. At least 1 week was allowed for animals to habituate to the new holding facility before surgery was performed. Post-surgery, animals were singly housed with additional bedding and cardboard tubes in high-roofed cages that allowed unconstrained head movement with cranial implants.

### Implantation of recording array

Custom-built adjustable tetrode (twisted 12.7μm nichrome wire, Kanthal, gold-plated to 250-300kΩ at 1kHz) microdrives were implanted under isoflurane anesthesia using aseptic technique and perioperative analgesia (Buprenorphine, 0.02mg/kg s.c.). Craniotomies of diameter 1-1.5 mm were made over dorsal CA1 (AP -4.2mm, ML 3.0mm from Bregma). Implants were fixed to the skull using stainless steel screws (M1.4 × 2 mm, Newstar Fastenings) and Gentamicin bone cement (DePuy). Tetrode positions were adjusted over the course of 2–3 weeks after surgery.

Tetrode signals were amplified by headstages (HS-36, Neuralynx, MT, USA) and relayed via fine-wire tethers to a Digital Lynx system (Neuralynx), which sampled thresholded extracellular action potentials at 32 kHz (filtered at 600-6000Hz) and continuous local field potentials (LFP) at 2kHz (filtered at 0.1-475Hz) using the Cheetah software package (Neuralynx) running on a desktop PC.

### Training

Once post-surgery body weight had stabilized, rats were placed on a regulated feeding regimen to maintain body weight at 85-90% of free-feeding levels. The rats were trained to perform a spatial memory-based decision-making task on an end-to-end T-maze, as described in Jones and Wilson [37] and illustrated in Figure 1a. Maze dimensions were 170 x 130 cm. Training occurred during the light phase at a similar time each day. *Habituation*: Rats were placed in the maze for 20–30 minutes without any boundaries in place. Rewards were provided at every visit to a reward zone. After 2 days, rats advanced to the next stage. *Guided trials*: Rats ran a series of ‘guided’ trials. Each trial consisted of a run from a reward point at one side of the maze, via the long central arm to a reward point at the opposite side. At the starting end of the maze, the opposite arm was blocked off with a barrier, guiding the rat onto the central arm. At the distal end of the central arm, a barrier blocking one of the arms (pseudorandomly selected) guided the rat to the end of the unobstructed arm, where a reward (0.1 ml of 20% sucrose solution in water) was delivered remotely through tubing connecting reward wells to syringe pumps located in the adjacent room, where the experimenter sat. Only one running trajectory was possible in each guided trial. In one training session, a rat was allowed to perform up to 40 trials. After a minimum of 2 days of at least 20 trials, rats advanced to the next training stage. *Full Task*: Rats performed a series of guided trials, interleaved with ‘choice’ trials. Choice trials differed from guided trials in that there was no barrier in place at the far end of the central arm, requiring the rat to choose a turn direction. The correct turn direction was the same direction that the rat had initially turned when entering the central arm. If the rat chose correctly, a reward was delivered at the end of the arm. If the rat chose incorrectly, it was placed back at the start and allowed to undertake the trial again, until the correct choice was made. All guided trials began at the ‘C’ end of the maze and ended at the ‘G’ end, while the interleaved choice trials ran in the opposite direction. Rats were allowed to perform up to 40/40 guided/choice trials. Learning of the task rule was assessed by the percentage of correct choices made (>70% correct trials over at least 3 consecutive days). In this manuscript, we analyzed recordings from four days (for a total of 24 sessions) after the performance criterion was reached.

### Histology

At the end of each experiment, the rat was deeply anesthetized with intraperitoneal sodium pentobarbital and a small electrolytic was lesion made at the tip of each tetrode (positive current of 0.3 mA for 10s). After lesions had been made on each tetrode, the rat was perfused transcardially with 0.9% saline and then 4% paraformaldehyde / 0.9% saline solution. The brain was post-fixed, transferred to a cold 30% sucrose solution for cryoprotection and cut in 50µm coronal sections on a freezing microtome. Lesion locations were compared against the corresponding sections in the Rat Brain Atlas [91] in order to determine the tetrode recording sites.

### Spike sorting and cell selection

Spikes were sorted semi-automatically on the basis of waveform characteristics (waveform energy and first principal component) using KlustaKwik (K.D. Harris, http://klustakwik.sourceforge.net/), followed by manual refinement of cluster boundaries with the MClust package for MATLAB (A.D. Redish, http://redishlab.neuroscience.umn.edu). After clustering, only units with mean spike peak amplitude of > 50µV, isolation distances of ≥15 [92] and <1% of interspike intervals (ISIs) below 2 ms were retained for further analysis. Our analysis focused on putative place cells. To select putative place cells we restrict to units with firing rate between 0.2 Hz and 4 Hz and with spatial information above 0.5 bit/s on the maze [39]. Because of the dependence introduced in the data by pooling units from multiple sessions of the same animals, when appropriate we perform statistical tests by generalized linear mixed-effects models, where we explicitly account for session and rat identity.

### Cell assembly detection

Cell assemblies were identified with the unsupervised machine learning algorithm for Cell Assembly Detection (CAD) [40] (algorithm available at https://github.com/DurstewitzLab/Cell-Assembly-Detection). CAD detects recurrent activity patterns of arbitrary structure and temporal precision in multivariate time series. The algorithm is based on a recursive agglomeration scheme, at each step of which it detects and tests assemblies of progressively larger size. As the first step of the algorithm, CAD scans the activity of all possible pairs of units in the recorded set to select the most common shared (potentially lagged) activation pattern, and test if this reoccurs more frequently than expected by chance. If the test is significant, a new time series is created to reflect the activation of the detected assembly-pattern. In the second step of the algorithm, the new assembly activation time series are tested in turn including the remaining recorded units. If the test is significant the unit becomes part of the assembly and a new assembly activation time series is generated. The algorithm stops when no more units can be added to the assemblies detected in the previous agglomeration step (see [40] for a more detailed description of the algorithm). It follows that the hypothetical inclusion of supplementary units beyond those actually recorded would affect the number and size of the detected assemblies but it would not alter the coordination patterns already detected in the existing set. In this manuscript, the detection of paired assemblies was performed stopping the agglomeration at the initial pairwise step, while full-size assemblies were detected letting the algorithm agglomerate until completion.

To uncover the temporal scales most represented in the hippocampal spike trains we ran CAD on a broad spectrum of temporal resolutions sampled with a logarithmic scale in the interval [0.005 - 5.0] sec. CAD tests multiple temporal resolutions and if the same sets of units coordinate at multiple timescales the algorithm will return all of them. This analysis revealed the presence of two distinct timescales: one between 0.005 and 0.06 sec (spike-assemblies) and a second between 0.07 and 5.0 sec (rate-assemblies). To compare the extent of phase-locking and phase shift-coding within the two assembly groups we repeated the assembly detection separately for the two time windows using CAD*opti* [93], [94] (algorithm available at https://github.com/DurstewitzLab/CADopti). After testing multiple temporal resolutions, CAD*opti* selects and returns the timescale at which each assembly has been detected with lowest p-value. Thus, each assembly was unique within each window but could be detected in both time windows. This pruning procedure allowed a fair comparison between the two timescales, without the distortion given by considering as independent assemblies the same set of units detected at neighboring temporal resolutions. Finally, we want to note that the detected assemblies cannot result from the detection of spike sorting mistakes. In such a case, in fact, assemblies would have been detected at the highest temporal precision (binning of 0.0058 sec), which is not the case as shown in Figure 1b.

For all the analysis of this manuscript the reference lag was set at 2. Tested bin sizes: [0.0058, 0.007, 0.009, 0.011, 0.014, 0.018, 0.022, 0.028, 0.035, 0.044, 0.055, 0.07, 0.09, 0.11, 0.14, 0.17, 0.21, 0.27, 0.33, 0.42, 0.52, 0.65, 0.82, 1.0, 1.3, 1.6, 2.0, 2.5, 3.2, 4.0, 5.0] and respective maximal lag: [4, 5, 5, 5, 5, 4, 4, 4, 3, 3, 3, 2, 2, 2, 2, 2, 1, 1, 1, 1, 1, 1, 1, 1, 1, 1, 1, 1, 1, 1, 1].

Values for tested bin sizes were selected based on previous experience in assembly analysis [40], [93], [94]. The lower limit was chosen because in CA1 we found practically no assemblies at coordination precision higher than 5 ms [40]. The upper limit was chosen to cover in about one bin the time needed by the rat to cross a place field.

### Computation of chance level for the probability of spike-assemblies being detected as rate-assemblies as well

Spike-assembly pairs had a probability of *p* = 0.9 to be also detected as rate-assembly pairs. We tested if the obtained value is above chance by bootstrap. To this aim, we considered all possible pairs of units present in the recorded sessions and randomly selected an amount equal to the number of the detected rate-assembly. We then computed the probability of the detected spike-assembly pairs to be part of this subset *p*^*i*^_*boot*_. We repeated the random sampling 10^5^ times and averaged the result to obtain the chance level 〈*p*_*boot*_〉 = 0.6. The p-value corresponds to the fraction of bootstrap sampling with *p* < *p*^*i*^_*boot*_, which, in the specific bootstrap set, never occurred.

### Instantaneous firing rate

Instantaneous firing rates were computed by convolving unit spikes with a Gaussian kernel with a kernel size of half of the unit’s mean inter-spike-interval.

### Phase extraction and theta power

As first step, we made sure that the spectrogram of all recorded LFPs peaked in the theta frequency band. Then, to obtain the phase of spikes we bandpass filtered the LFP between 4Hz and 10Hz (LFPθ) and computed the angle of the Hilbert transformation of LFPθ at the time of each spike.

Since recent studies have shown that phase coding can occur also in the presence of a lower power or irregular amplitude of the theta oscillation [95], [96], the analyses reported in the main figures of the manuscript are performed with maximal sample size, including all spikes without restriction on theta power. However, to make sure that the reported results were not affected by this choice, we reproduced the most important including only spikes fired in epochs of high theta power (Supplementary Fig. 9 and 11). High theta power epochs were defined as periods in which the envelope of the LFPθ amplitude surpassed one *σ*(LFPθ). High amplitude, saturating movement artifacts were removed from LFP by excluding periods with LFPθ > 2 ∗ *σ*(LFPθ). This was done before computing the threshold of *σ*(LFPθ) used to define high power periods.

### Detection of SWR

SWR detection was based on the ‘Sleepwalker’ MATLAB toolbox (https://gitlab.com/ubartsch/sleepwalker). In brief, LFPs were down-sampled to 1000 Hz and 50 Hz notch filtered prior to band-pass filtering (using least squares filters) between 120-220 Hz. Candidate ripple events were identified based on threshold crossings in the z-scored 120- 220 Hz power, using a threshold of 3.5 x SD of the signal. Start and finish times of the ripple were calculated based on 2 x SD of 120-220 Hz power. Representative samples of individual and averaged events were then visually inspected. Ripples were rejected if they were shorter than 50 ms or longer than 500 ms; if gaps of less than 50 ms occurred between events, they were treated as a single ripple. Ripple start/end timestamps were used to exclude SWR-associated spiking for some analyses (**Supplementary** Fig. 4, 5, 9, 11).

### Phase-locking test

Phase modulation of spike sets was tested with the Hodges-Ajne test for non-uniformity [97], [98] that, unlike the more common Rayleigh test, does not assume unimodality in deviation from uniformity. The test was performed on sets bigger than 50 samples, limit imposed by the approximations performed in the test. Significance was established with an alpha value of 0.05 and Benjamini-Hochberg correction for multiple comparisons on all tests performed.

### Phase-locking of assembly-spikes vs. unit-spikes

This test compares the strength of phase-locking of two sets of phase values. In particular, we tested via bootstrap if assembly activations elicited spikes with higher phase-locking than those overall fired by the unit. For each unit, we collected the phase of the *n* spikes fired in correspondence with all activations of one assembly and produced 1000 replica sets composed of the phase of *n* spikes randomly selected among all spikes of the same unit. For each set, we computed the length *R* of the mean resultant vector. P-values were established by counting the fraction of replica-sets with *R*_*rep*._ > *R*_*orig*._. Significance was established with an alpha value of 0.05 and Benjamini-Hochberg correction for multiple comparisons on all tests performed.

### Change in phase preference for spikes fired in different assemblies

This test assesses if two sets of phase values have the same median phase. For each unit taking part in multiple assemblies, we divided into separate sets the spikes fired in different assembly-pairs. Sets not phase-locked to the ongoing theta oscillation or with less than 50 spikes (limitation imposed to perform the phase-locking test) were discarded. To test the null hypothesis that for a same unit each phase-set had equal median, we performed a multi-sample nonparametric test, circular analog to the Kruskal-Wallis test [97], [99]. Significance was established with an alpha value of 0.05 and Benjamini-Hochberg correction for multiple comparisons on all tests performed. Finally, we tested whether the number of units recorded in each session correlated with the fraction of units per session changing phase when firing in different assemblies. We found no significant correlation (Spearman’s correlation *r*_*s*_(34) = 0.25, *p* − *value* = 0.14).

### Trial categories

We divided trials into 6 categories according to the specific task required by each trial. Trials were first divided into *choice trials*, when the animal had to choose if to turn left or right on the basis of its position at the beginning of the trial, and *guided trials*, when turns were forced by the set-up. Both categories were then further divided according to the two turns performed entering and leaving the central arm of the maze: *left-left* / *left-right* / *right-left* / *right-right*.

### Isolation of place fields

Place fields were established only for units with spatial information above 0.5 (*place cells*) and with phase modulated spikes. Spikes of each unit were divided into different clusters (place fields) on the basis of their place of firing. Place fields are often identified as a region of connected bins in a unit’s rate map that surpasses a fixed threshold in firing rate. For example, for a maze comparable to the one used in this manuscript, Wirtshafter and Wilson [100] use a threshold of a standard deviation over the mean firing rate of the unit. In units with multiple place fields, such a high threshold leads to a selection among the fields, at the expense of those with lower rates which remain undetected (c.f. Supplementary Fig. 7). While this is typically not an issue, this study aims to investigate the changes in phase and rate a unit exhibits across different place fields and trial types. Therefore, it is here important to capture a larger variety of fields. We thus explored different techniques to detect place fields and tested the robustness of the analyses in Fig. 5b and 5d.

The compared techniques were:

### Rate Map Wirtshafter and Wilson (RM WW)

In Wirtshafter and Wilson [100] place fields are detected by: 1) computing a rate map of firing per occupancy binned with a 2 cm grid and smoothed with a 10 cm standard deviation Gaussian kernel. Only epochs in which the rat moved faster than 12 cm/s were included; 2) thresholding the rate map at θ^1^_*RMWW*_ = μ(*fr*) + *σ*(*fr*), with μ(*fr*) and *σ*(*fr*) rate mean and standard deviation, respectively; 3) only fields with at least one bin with rate above θ^2^ = μ(*fr*) + 2 · *σ*(*fr*) were selected; 4) fields of length less than 15 cm were discarded.

### Rate Map (RM)

To detect also place fields with lower firing rates we repeated the procedure described in ‘RM WW’ but using a θ^1^_*RM*_ = 0.7 · μ(*fr*) and omitting point 3.

We also included methods that aimed to group spikes according to the density of their spatial clustering:

### Rate Map + GMM (RM + GMM)

We modeled the place fields of a unit with a Gaussian Mixture model. In the first step, we proceeded as described in ‘RM’ but with a harsher threshold of *θ*^1^_*RMGMM*_ = μ(*fr*). This allowed us to establish the overall number of place fields and obtain a first estimation of cluster memberships. In the second step, this first clustering was then used as initial condition for the estimation of a Gaussian mixture model (function *fitgmdist*, matlab). To train the model we only used the spikes identified as place field members in the first step. Once obtained the model we used it to cluster (function *cluster*, Matlab) all spikes fired by the unit. Spikes with low probability of being part of any of the identified clusters (i.e. with logarithm of the estimated probability density function smaller than *logpdf* < −20) were discarded as not assigned to any of the modeled Gaussians (place fields).

### DBSCAN

Spikes were clustered with a density-based spatial clustering algorithm (DBSCAN [101], Matlab function *dbscan* with parameters ε = 0.05 and *MinPts* = 15). Spikes identified as outliers by the algorithm were discarded.

### DBSCAN + GMM

We proceeded as in ‘RM + GMM’ but used the output of ‘DBSCAN’ as the initial place field estimate.

Common to all tested place field detection methods, only spikes fired when the rat moved faster than 12 cm/s were included.

Supplementary Fig. 7 shows a comparison between the outputs of the different place field detection methods. Key analyses were repeated on unit’s activity parsed by place field computed with different detection methods (Supplementary Fig. 8, 9, 10, 11, and text).

### Classifier

For each place field we trained a support vector machine (SVM) classifier with linear kernel (slack variables minimized with L1 norm and box constraint = 1) to divide trials according to their trial type (the trial types categories here considered were: correct choice left, correct choice right, forced left, forced right, forced switch right-left, forced switch left-right). For every two types of trials, we build 3 SVM models: one based on the spike’s phase, one on the spike’s instantaneous firing rate, and one on both phase and instantaneous firing rate. Spike phases are a circular quantity and cannot be used directly to train the SVM. Thus, phase information was passed to the classifier as [cos(*θ*_*t*_), sin(*θ*_*t*_)], where *θ*_*t*_ is the phase relative to the theta band of the LFP of the spike fired at time *t*. The classifier accuracy was computed with a 50-fold cross-validation, to avoid overfitting. Significance was established via bootstrap. Bootstrapped samples were created by shuffling the trial labels. Since this step removes not only the spike-label association but also any autocorrelation in the label time series (which might affect the accuracy when performing block cross-validation), for a fair comparison, we jointly shuffled the order of the spike-label elements when training the SVM classifier on the original set. The bootstrap procedure was repeated 500 times and p-values were assigned by counting the fraction of bootstrap sets with an accuracy higher or equal to the original set.

### Phase precession units

We tested phase precession for each place field of each unit. Precession was assessed separately for each trial type by computing the circular-linear correlation [97], [98] between the unit phase and the position of the animal along the linearized trials-specific path when the spike was fired.

### Changes in lag of activation between units along theta cycles

To assess if pairs of units significantly changed their relative lag of activation during different trial types, we first computed the maximal cross-correlation lag of the two units in each trial. Cross-correlation was computed with a 0.02 sec binning and, to remain within a theta cycle, within the [-3, 3] bin window (in Figure 6e we chose a larger window exclusively for visualization purposes). Once obtained the maximal correlation lag per trial, we divided the trials according to their trial type and tested for a change in lag with a two-sided Wilcoxon signed rank test. Since single units had different phase preferences in different place fields, testing was performed separately for each unit place field.

### AdEx model and assembly recruitment

Neuronal activity was simulated by an Adaptive Exponential Integrate-and-Fire model [51]. In AdEx models, the evolution of the neuron membrane potential *V* and adaptation current *w* is defined by the equations:

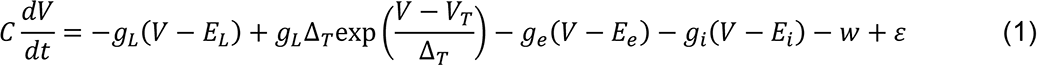

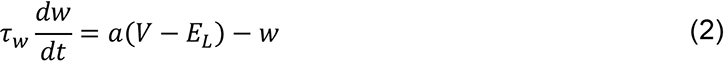

With membrane capacitance *C*, leak conductance *g*_*L*_, threshold slope factor Δ_*T*_, resting potential *E*_*L*_, threshold potential *V*_*t*_, adaptation time constant τ_*w*_, and subthreshold adaptation *a*. Noise in the evolution of the membrane potential was introduced through the parameter ε∼*N*(0,1.6 10^−19^). Synaptic currents were modulated through the time-dependent excitatory and inhibitory conductances *g*_*e*_(*t*) and *g*_*i*_(*t*), and the excitatory and inhibitory current reversals *E*_*e*_ and *E*_*i*_, respectively. At every time *t̅* the membrane potential reached 0 mV, an action potential was fired and *V*(^*t̅*^) was set to *V*_*t̅*_. Afterward, both the membrane potential and the adaptation current were reset to *V*(^*t̅*^ + 1) = *V*_*r*_ and *w*(*t̅* + 1) = *w*(*t̅*) + *b*, respectively.

To reflect the presence of theta oscillations, we modulated both excitatory and inhibitory conductances at *θ* = 7 Hz with a relative offset of π rad. The excitatory conductance was then further modulated by a gaussian-shaped depolarization to mimic transient assembly activation

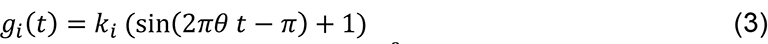

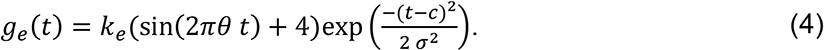

with *c* = 2.5 *sec* and *σ* = 0.4 *sec* center and standard deviation of the Gaussian, respectively. The activation of an assembly provided to its composing unit *n* an assembly-specific degree of depolarization *k*^*n*^_*e*_. We simulated 2 assemblies composed of 3 units. In the first assembly *k*^1^_*e*_ = 2.7 *nS*, *k*^2^_*e*_ = 1.7 *nS* and *k*^1^_*e*_ = 0.7 *nS*. In the second assembly *k*^1^_*e*_ = 1.0 *nS*, *k*^2^_*e*_ = 2.0 *nS* and *k*^1^_*e*_ = 3.0 *nS*. Average inhibitory conductance was set at *k*_*i*_ = 17 *nS* for all units and all assemblies.

To formally evaluate the fine temporal coordination of network units induced by the recruitment of different context-specific cell assemblies, we concatenated 400 retrievals of each of the two modeled assemblies and ran CAD*opti* on the spike time series so obtained. We performed and analyzed 100 simulations generated with different noise realizations. CAD*opti* parameters: reference lag = 2; bin sizes: [0.0058, 0.007, 0.009, 0.011, 0.014, 0.018, 0.022, 0.028, 0.035, 0.044, 0.055]; maximal lag: [18, 21, 22, 22, 22, 20, 18, 17, 15, 13, 11].

## Supplementary Information

### Supplementary figures

**Supplementary Fig. 1.**
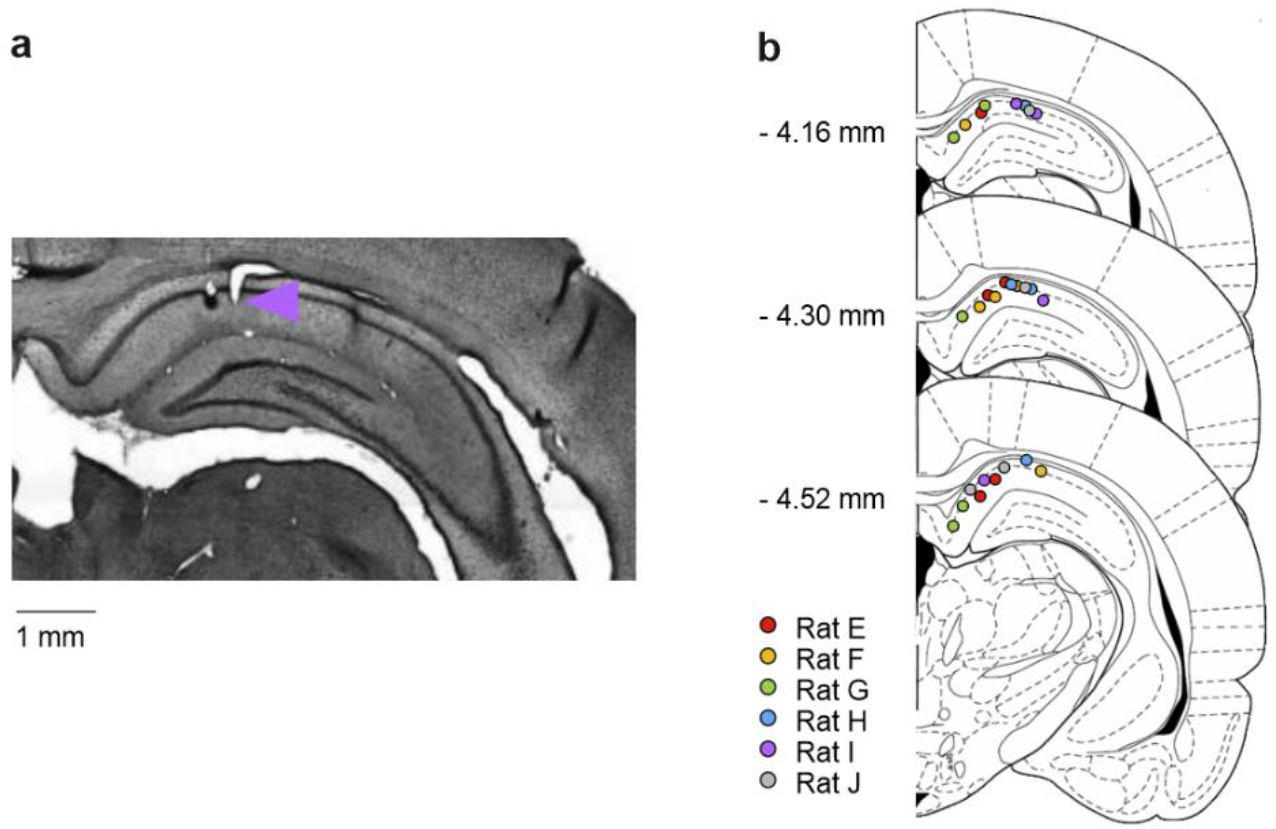
Histology. (a) Coronal section showing histological confirmation of the tetrode placement (lesion site marked with arrowhead) in the CA1 of one rat; (b) Dorsal CA1 tetrode placement for all recorded rats.

**Supplementary Fig. 2.**
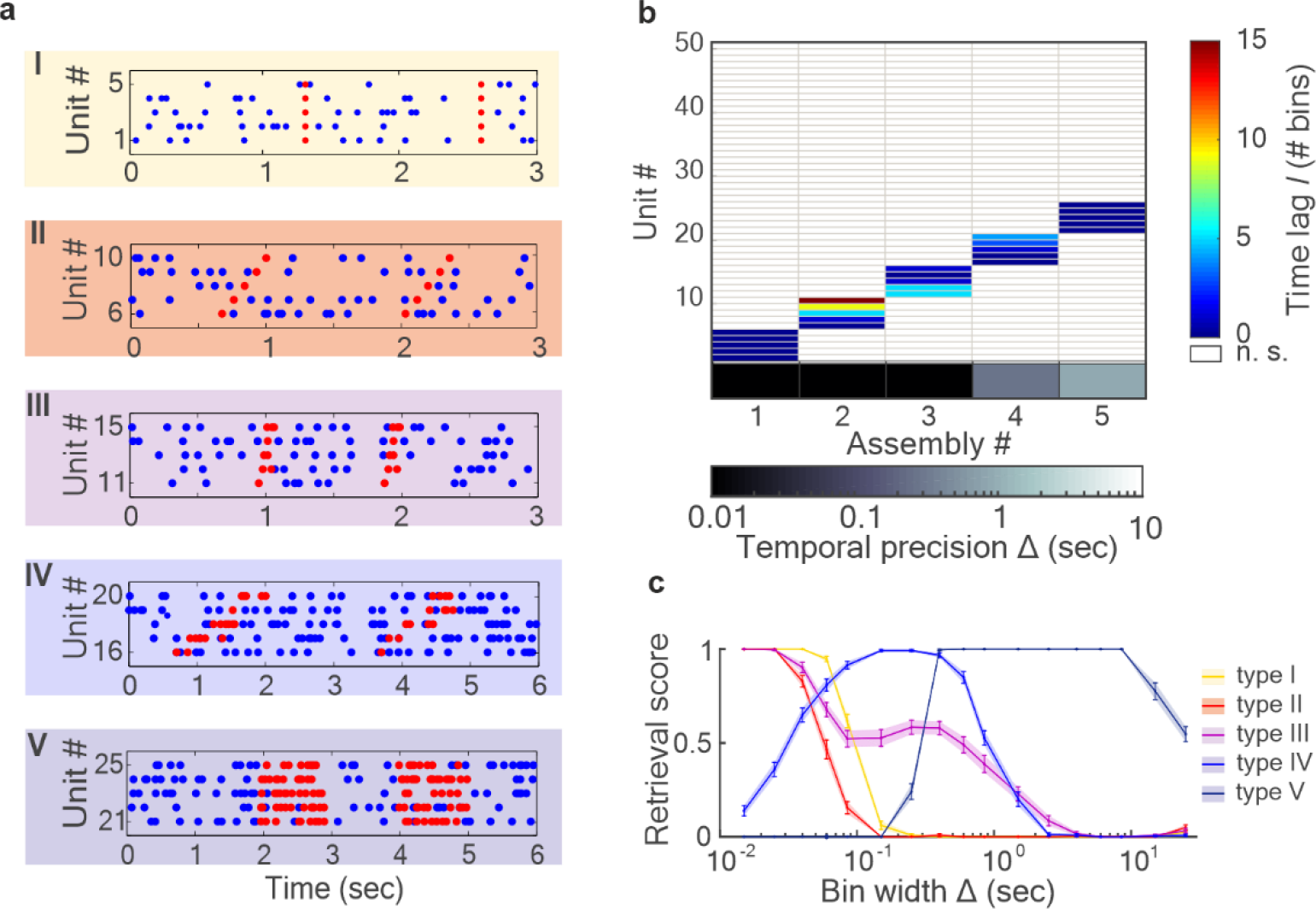
Cell assembly detection with CAD. Hippocampal assemblies were detected by applying the method for Cell Assembly Detection (CAD) presented in [40]. CAD is able to detect assemblies in an unsupervised fashion returning for each assembly the identity of its composing units, the activation pattern, and the temporal resolution of the unit coordination. To exemplify the type of information returned by CAD, we here report the algorithm performance when applied to a set of 50 simulated units containing 5 distinct assemblies of 5 units each. To showcase the wide range of CAD sensitivity, the simulated assemblies differed both in activation pattern and temporal resolution. (a) Rasteplots of 25 of the 50 simulated units (in blue), showing the activation of the five simulated assemblies (in red): type I – highly precise lag-0 synchronization; type II – highly precise sequential pattern; type III – highly precise spike-time pattern without clear sequential structure; type IV – rate pattern with sequential pattern; V – synchronous rate increase. Units from 26 to 50 were not included in any assembly. (b) Assembly-assignment matrix showing the output of CAD. Each detected assembly corresponds to a column of the assembly-assignment matrix. Colored units (rows) have been identified as part of the assembly. The color indicates the lag between the activation of the unit and that of the first unit active within the assembly. Along the abscissa, in grey-scale, the temporal resolution of the detected assembly. (c) Fraction of correctly assigned units (retrieval score) as a function of the temporal resolution (bin size) at which the algorithm scans the spike train. Data averaged across 70 independent runs, error bars = SEM. Thanks to the non-stationarity correction implemented in the algorithm (see [40]), CAD detects assemblies only at their characteristic temporal resolution. Thus, precise assemblies of type I and II were only detected at very small bin sizes, while rate assemblies of type IV and V only at bigger bin sizes. Interestingly, the type III assembly was detected both at small and larger bin sizes because of the precise timing in the relative activation of assembly units, and the change in firing rate produced by its extended activation pattern. Figure taken from ref. [40] under CC-BY 4.0 license.

**Supplementary Fig. 3.**
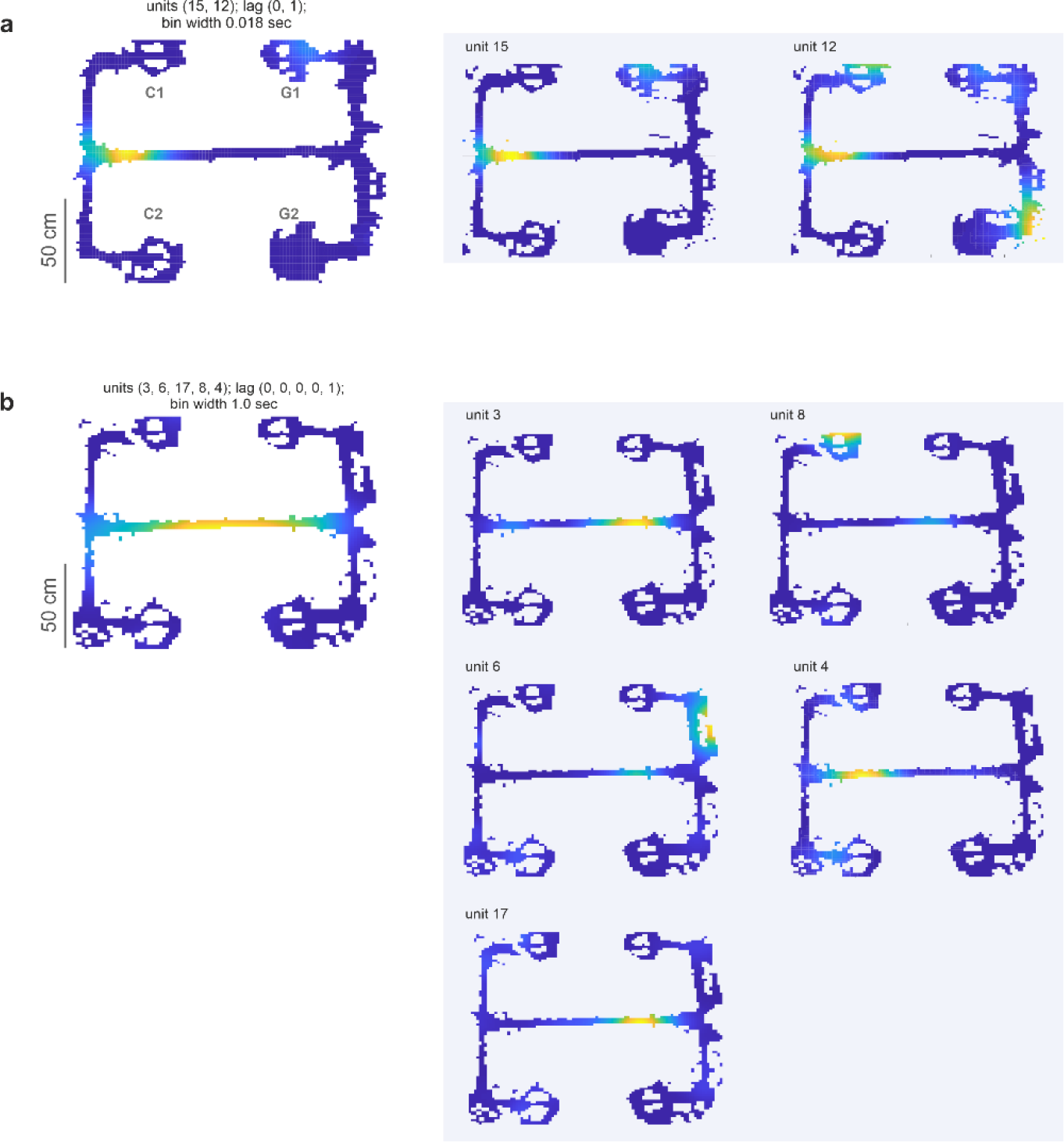
Activation maps of assemblies and their composing units. Example of activation maps of a spike- (a) and a rate- (b) assembly and of its composing units. While single units fired in multiple place fields throughout the maze, assembly activations were more selective.

**Supplementary Fig. 4.**
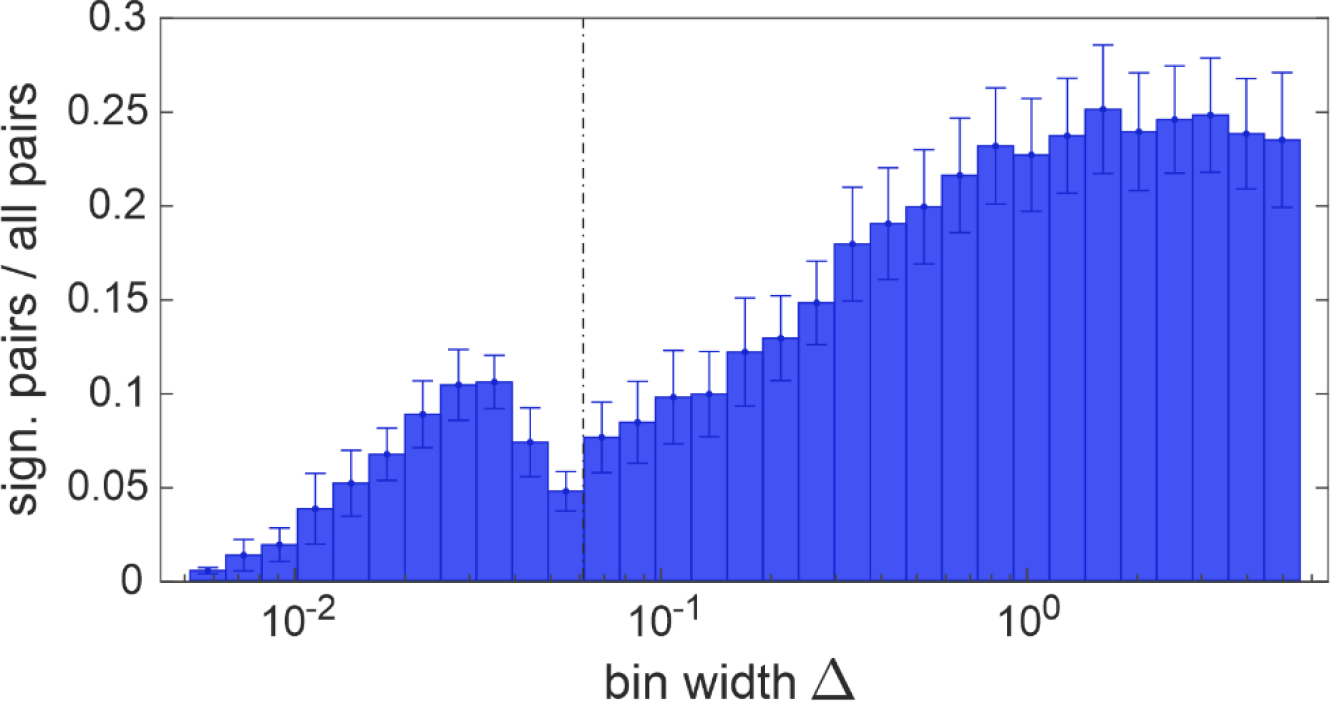
Temporal precision of CA1 assemblies detected outside of SWR epochs. Same as in Fig. 1b, but with assemblies detected on spike trains where epochs with SWR were excluded.

**Supplementary Fig. 5.**
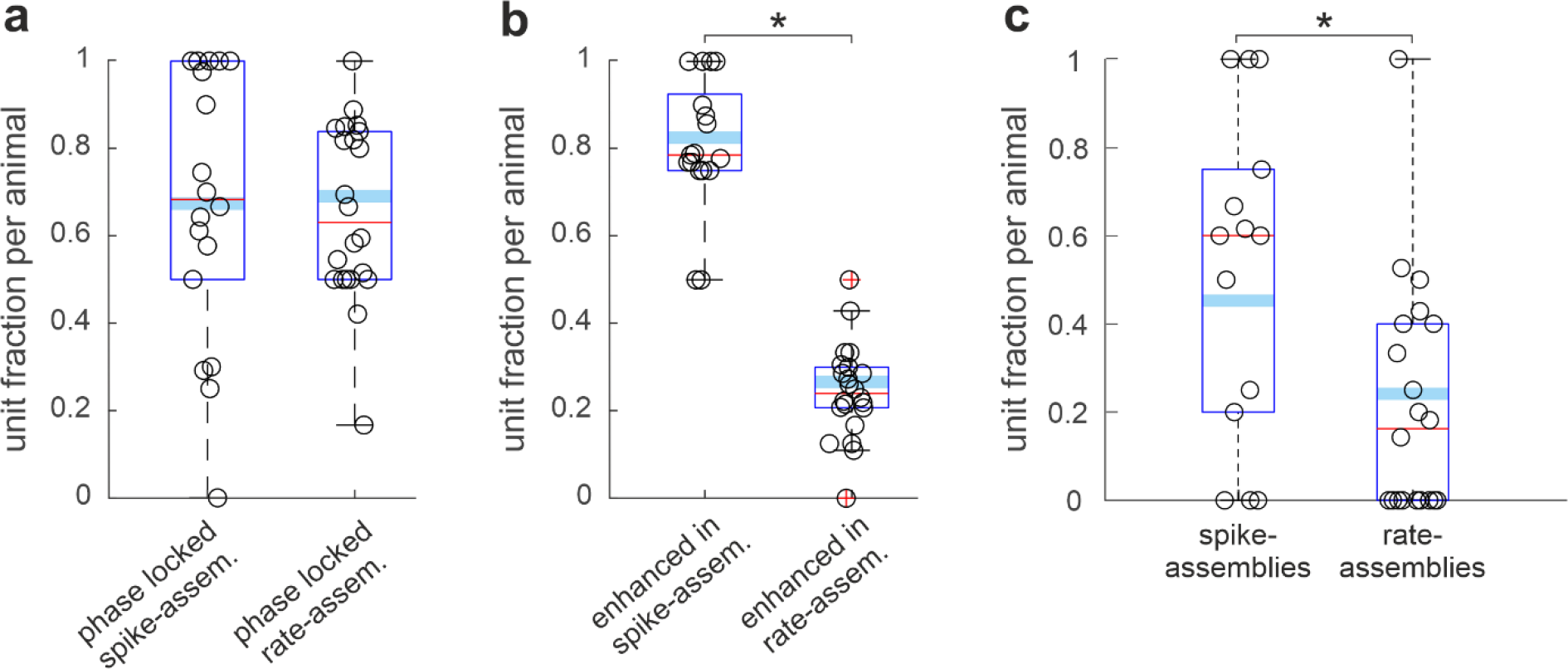
Theta-phase modulation of assemblies detected outside of SWR epochs. (a) Same as Fig. 2b, (b) same as Fig. 2c, (c) same as Fig. 3b but computed on assemblies detected excluding epochs with SWR. In all three panels we tested significance with generalized linear mixed-effects models to account for rat identity and recording session as covariates. The models and their output statistics are (a) generalized linear mixed-effects model of the probability of a unit to phase-lock when firing within an assembly according to the assembly type, spike- vs rate-assembly: F(1,1812) = 1.9, p = 0.17; (b) generalized linear mixed-effects model of the probability of a unit to increase phase modulation when firing within an assembly according to the assembly type, spike- vs rate-assembly: F(1,1301) = 178.4, p = 3.2 10^-38^; (c) generalized linear mixed-effects model of the probability of a unit to phase-shift when firing in different assemblies according to the assembly type, spike- vs rate-assembly: F(1,230) = 6.93, p = 9.0 10^-3^.

**Supplementary Fig. 6.**
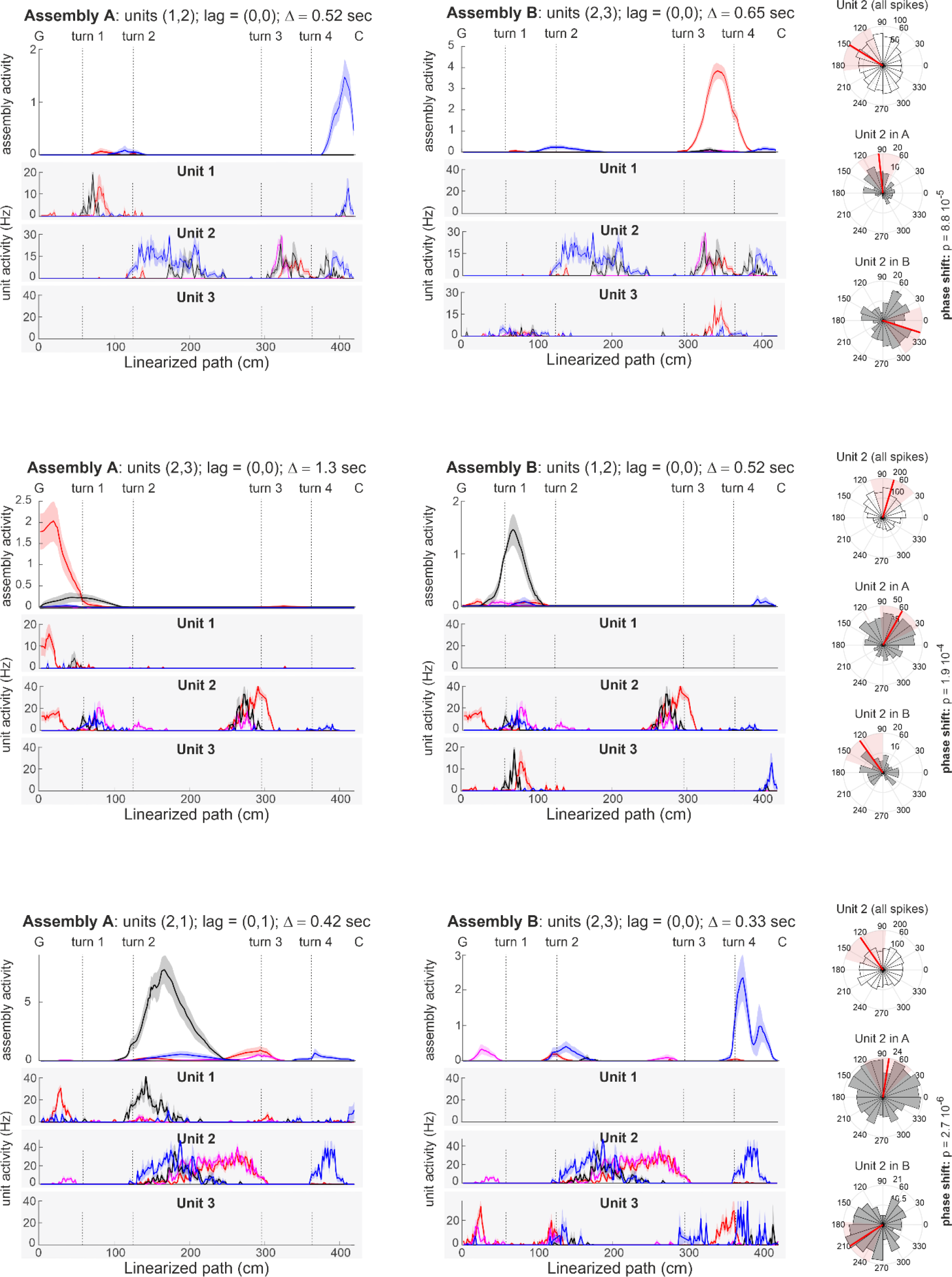
Changes in unit information coding when joining different assemblies. Same as Fig. 3c, three additional examples of units with changing phase preference when active in different assemblies.

**Supplementary Fig. 7.**
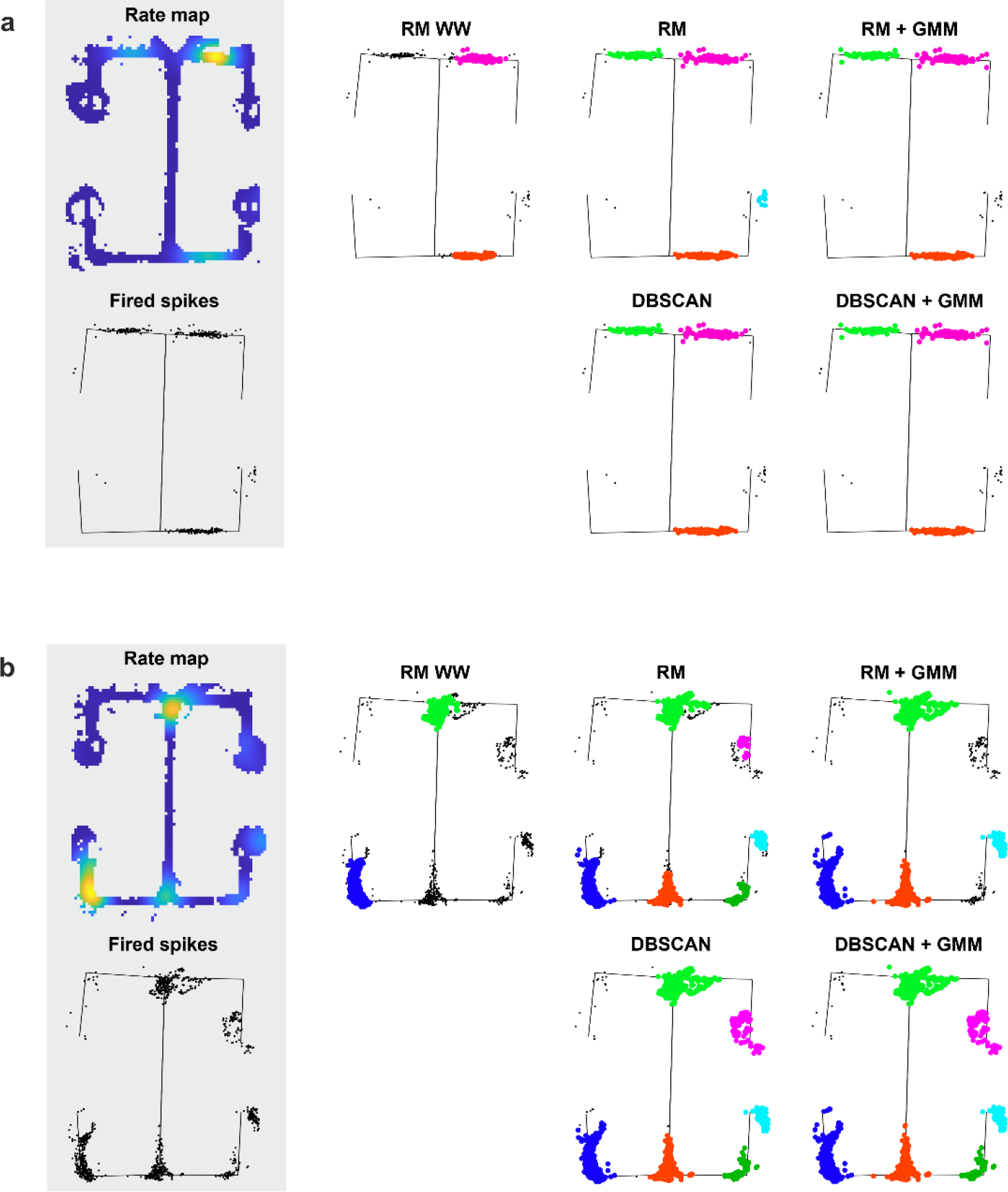
Method comparison for place field detection. We compared the output of 5 different place field detection methods: a thresholded rate map based detection algorithm presented in Wirtshafter and Wilson [100] (RM WW); a rate map based detection algorithm adapted from Wirtshafter and Wilson (RM); the RM algorithm combined with a Gaussian mixture model (RM + GMM); a density-based spatial clustering algorithm (DBSCAN); the DBSCAN algorithm combined with a Gaussian mixture model (DBSCAN + GMM) (see Methods for discussion and description of the place field detection algorithms). In (a) and (b) in the grey panel the rate map (top) and the position on the maze (bottom) of the recorded spikes of two example place cells. On the right, spike’s positions color-coded according to the place fields detected by the 5 different algorithms. Different colors corresponded to different place fields. As visible in the two examples, in units with multiple place fields, thresholding the rate map at one standard deviation above the average firing rate of the unit might result in the failure to detect those fields where the unit consistently becomes active at lower frequencies.

**Supplementary Fig. 8.**
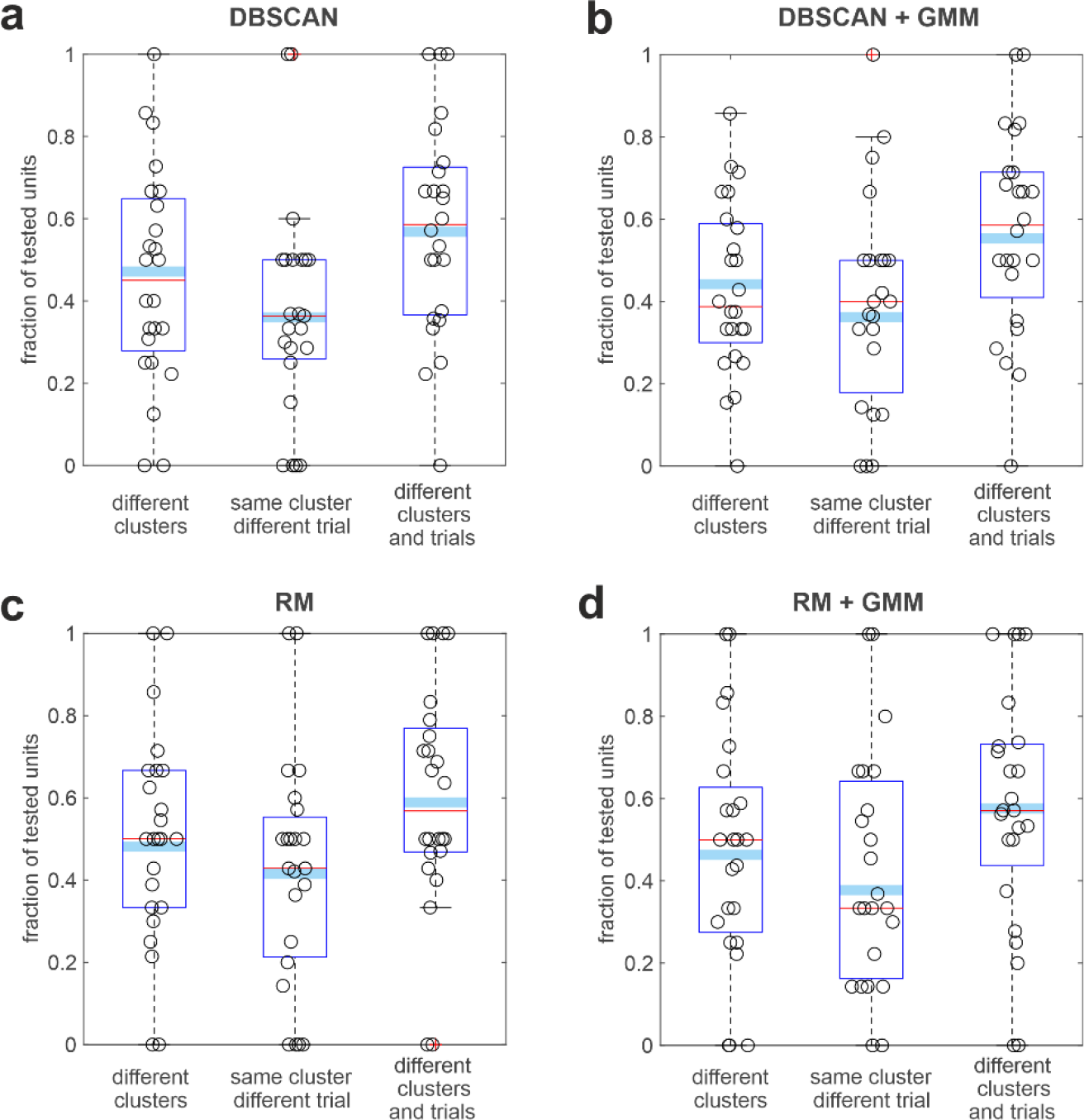
Fraction of units changing preferred theta-phase for different place fields and trial types. To probe the robustness of the results reported in Fig. 5b we performed the same analysis on sets of place fields identified with alternative techniques. From panel (a) to (d) we used place fields detected respectively by: a density-based spatial clustering algorithm (**DBSCAN**); the DBSCAN algorithm combined with a Gaussian mixture model (**DBSCAN + GMM**); thresholding the unit rate map (**RM**); the RM algorithm combined with a Gaussian mixture model (**RM + GMM**) (see Methods for discussion and description of the place field detection algorithms). Panel (b) is the same as Fig. 5b and reported here for comparison.

**Supplementary Fig. 9.**
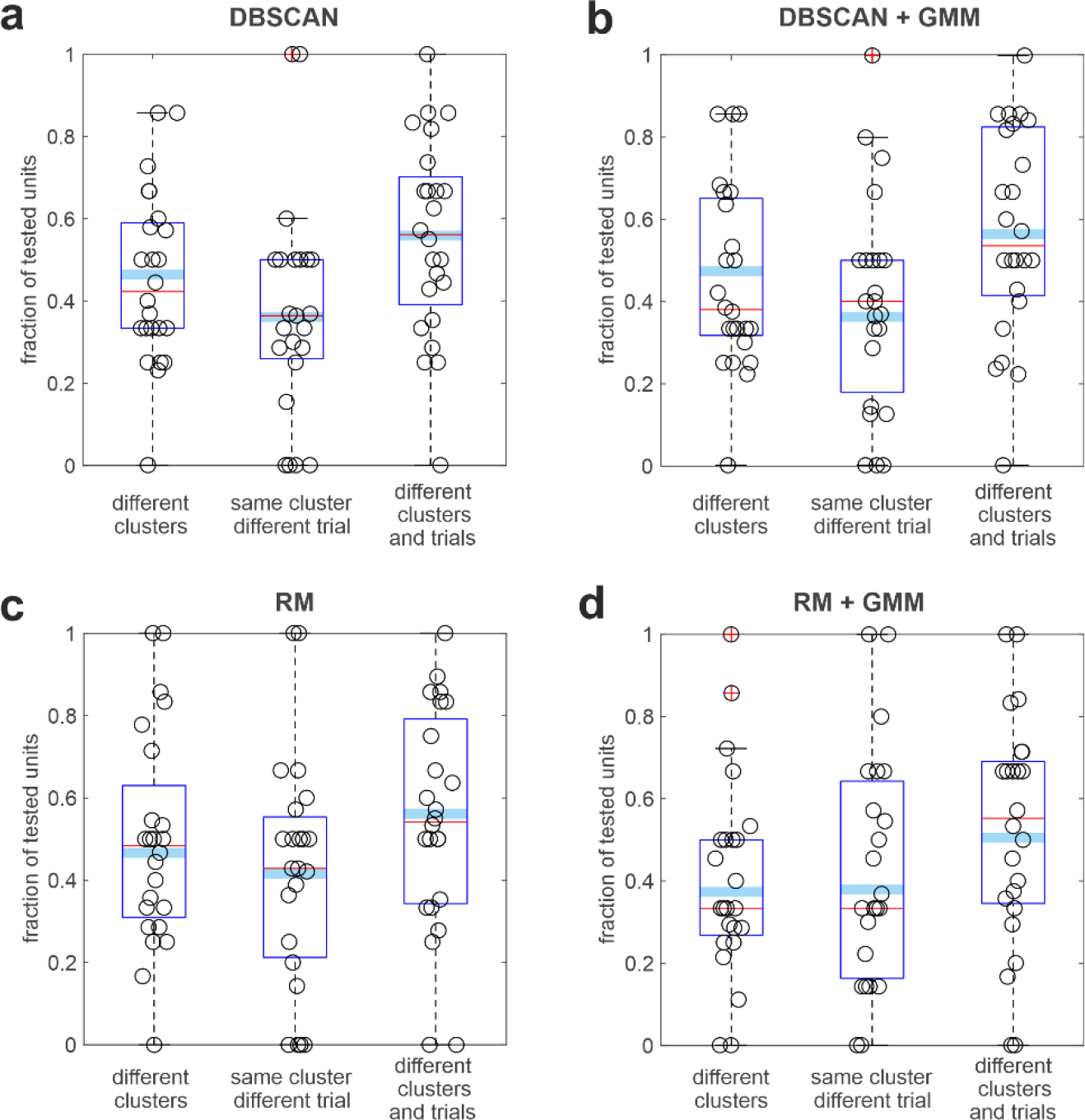
Fraction of units changing preferred theta-phase for different place fields and trial types (only high theta power and no SWR included). Same as Supplementary Fig. 8 but with analyses performed on spikes fired in epochs of high theta-power and outside of SWR epochs.

**Supplementary Fig. 10.**
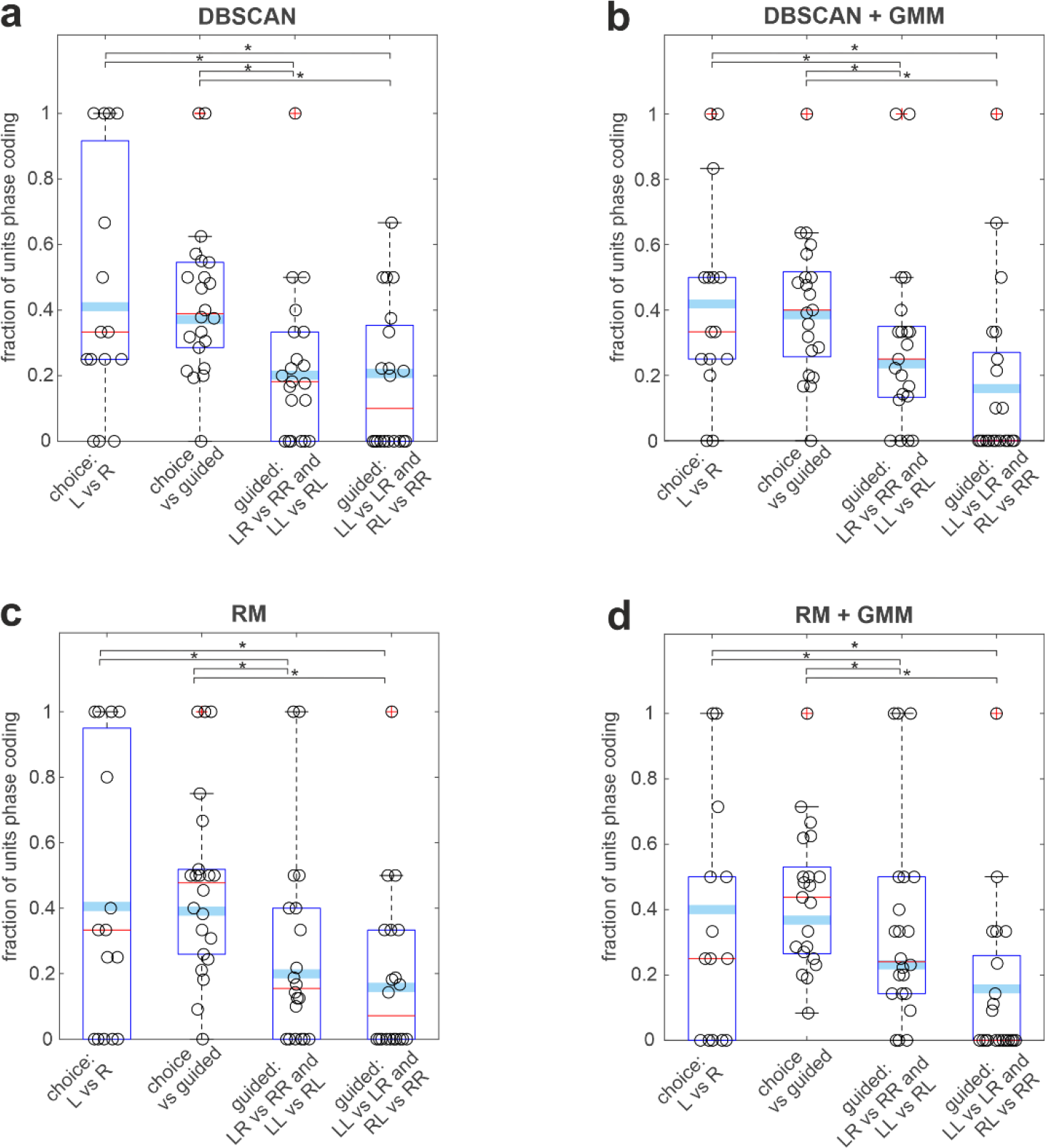
Fraction of units changing preferred theta-phase for different sets of trial types. To probe the robustness of the results reported in Fig. 5d we performed the same analysis on sets of place fields identified with alternative techniques. From panel (a) to (d) we used place fields detected respectively by: a density-based spatial clustering algorithm (DBSCAN); the DBSCAN algorithm combined with a Gaussian mixture model (DBSCAN + GMM); thresholding the unit rate map (RM); the RM algorithm combined with a Gaussian mixture model (RM + GMM) (see Methods for discussion and description of the place field detection algorithms). In all four panels we tested significance with a generalized linear mixed-effects model to account for rat identity and recording session as covariates. We modeled if a unit changed firing phase (binary 0/1 variable and logit link function) within the same place field according to the trial types. The resultant statistics was (a) global F-statistics F(4,613) = 14.4, p = 3.3 10^-11^; contrast tests between specific condition/bars (I vs II) F(1,613) = 0.2, p = 0.6; (I vs III) F(1,613) = 6.9, p = 8.6 10^-3^; (I vs IV) F(1,613) = 5.6, p = 1.8 10^-2^; (II vs III) F(1,613) = 12.8, p = 3.7 10^-4^; (II vs IV) F(1,613) = 8.5, p = 3.6 10^-3^; (III vs IV) F(1,613) = 0.0, p = 0.9; (b) global F-statistics F(4,641) = 10.2, p = 4.9 10^-8^; contrast tests between specific condition/bars (I vs II) F(1,641) = 0.2, p = 0.6; (I vs III) F(1,641) = 5.6, p = 1.8 10^-2^; (I vs IV) F(1,641) = 10.7, p = 1.2 10^-3^; (II vs III) F(1,641) = 9.8, p = 1.8 10^-3^; (II vs IV) F(1,641) = 16.4, p = 5.8 10^-5^; (III vs IV) F(1,641) = 1.9, p = 0.2; (c) global F-statistics F(4,597) = 9.3, p = 2.9 10^-7^; contrast tests between specific condition/bars (I vs II) F(1, 597) = 0.1, p = 0.7; (I vs III) F(1, 597) = 7.2, p = 7.7 10^-3^; (I vs IV) F(1, 597) = 9.5, p = 2.1 10^-3^; (II vs III) F(1, 597) = 14.8, p = 1.3 10^-4^; (II vs IV) F(1, 597) = 16.6, p = 5.1 10^-5^; (III vs IV) F(1, 597) = 0.6, p = 0.5; (d) global F-statistics F(4,631) = 9.3, p = 2.7 10^-7^; contrast tests between specific condition/bars (I vs II) F(1, 631) = 0.2, p = 0.7; (I vs III) F(1, 631) = 4.3, p = 3.9 10^-2^; (I vs IV) F(1, 631) = 9.1, p = 2.7 10^-3^; (II vs III) F(1, 631) = 8.1, p = 4.7 10^-3^; (II vs IV) F(1, 631) = 15.0, p = 1.2 10^-4^; (III vs IV) F(1, 631) = 2.0, p = 0.2. Panel b is the same as Fig. 5d and reported here for comparison.

**Supplementary Fig. 11.**
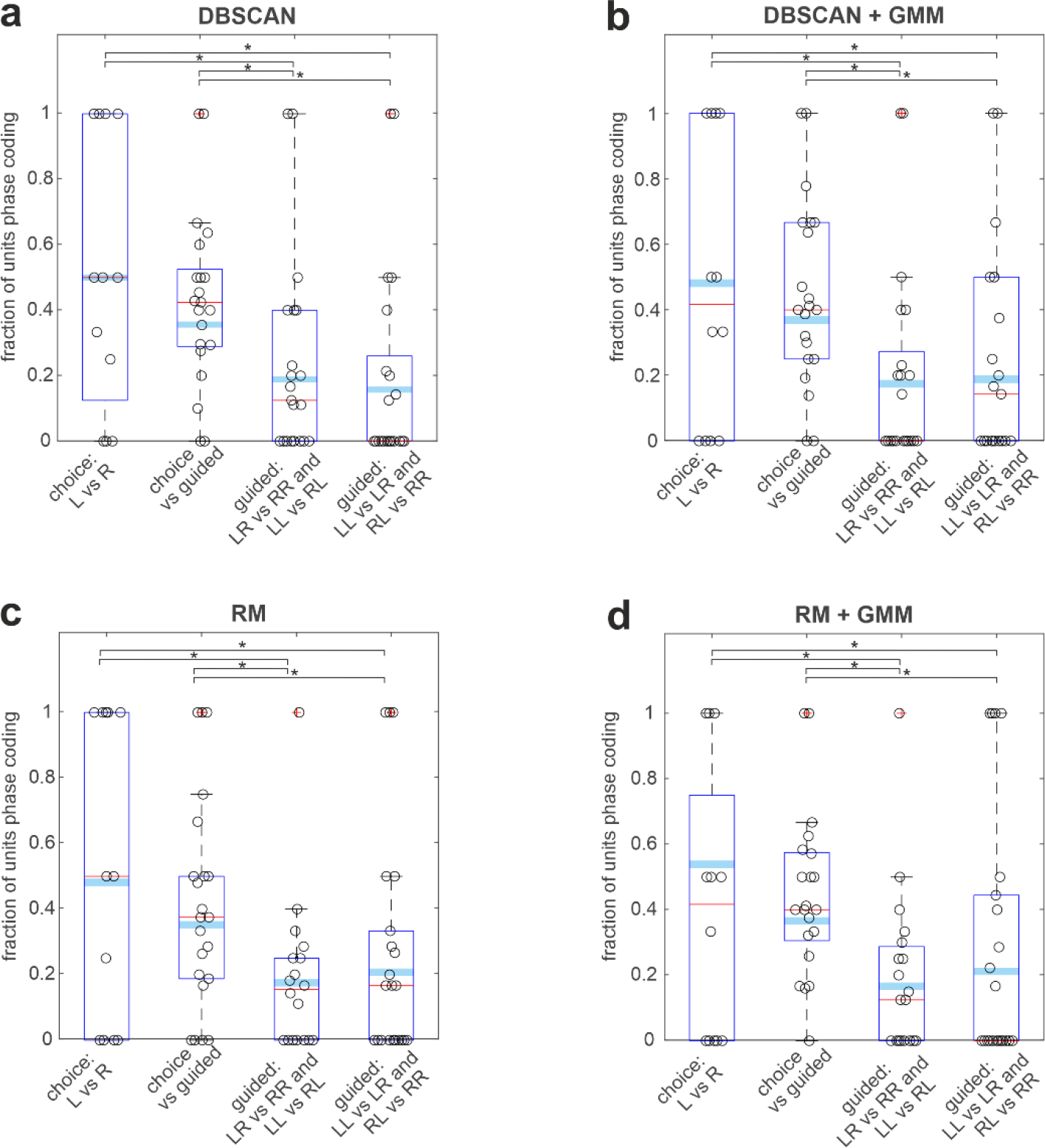
Fraction of units changing preferred theta-phase for different sets of trial types (only high theta power and no SWR included). Same as Fig. 5d and Supplementary Fig. 10 but with analyses performed on spikes fired in epochs of high theta-power and outside of SWR epochs. In the four panels the resultant test statistics was (a) global F-statistics F(4,514) = 10.0, p = 8.2 10^-8^; contrast tests between specific condition/bars (I vs II) F(1,514) = 2.2, p = 0.1; (I vs III) F(1,514) = 11.0, p = 1.0 10^-3^; (I vs IV) F(1,514) = 12.5, p = 4.5 10^-4^; (II vs III) F(1,514) = 11.5, p = 7.6 10^-4^; (II vs IV) F(1,514) = 12.6, p = 4.2 10^-4^; (III vs IV) F(1,514) = 0.3, p = 0.6; (b) global F-statistics F(4,515) = 8.2, p = 2.0 10^-6^; contrast tests between specific condition/bars (I vs II) F(1,515) = 1.5, p = 0.2; (I vs III) F(1,515) = 11.1, p = 9.4 10^-4^; (I vs IV) F(1,515) = 8.9, p = 3.0 10^-3^; (II vs III) F(1,515) = 14.0, p = 2.0 10^-4^; (II vs IV) F(1,515) = 9.6, p = 2.0 10^-3^; (III vs IV) F(1,515) = 0.1, p = 0.7; (c) global F-statistics F(4,507) = 11.7, p = 4.2 10^-9^; contrast tests between specific condition/bars (I vs II) F(1,507) = 2.6, p = 0.1; (I vs III) F(1,507) = 13.4, p = 2.8 10^-4^; (I vs IV) F(1,507) = 9.5, p = 2.1 10^-3^; (II vs III) F(1,507) = 13.7, p = 2.3 10^-4^; (II vs IV) F(1,507) = 7.6, p = 6.0 10^-3^; (III vs IV) F(1,507) = 0.5, p = 0.5; (d) global F-statistics F(4,472) = 7.7, p = 5.3 10^-6^; contrast tests between specific condition/bars (I vs II) F(1,472) = 0.2, p = 0.1; (I vs III) F(1,472) = 10.7, p = 1.1 10^-3^; (I vs IV) F(1,472) = 7.5, p = 6.5 10^-3^; (II vs III) F(1,472) = 10.2, p = 1.5 10^-3^; (II vs IV) F(1,472) = 5.3, p = 2.1 10^-2^; (III vs IV) F(1,472) = 0.4, p = 0.5. This control confirmed that excluding spikes fired at low theta power and during SWR did not change the observed modulation in theta phase according to the trial type of some CA1 units.

**Supplementary Table 1.** Parameters of the adaptive exponential integrate-and-fire model. Parameters used to produce the results presented in Fig. 6f.

| Parameter | Value |
| --- | --- |
| $C$ | 250 pF |
| $g_L$ | 10 nS |
| $E_L$ | -58 mV |
| $\Delta_T$ | 2 mV |
| $V_T$ | -50 mV |
| $E_e$ | 0 mV |
| $E_i$ | -75 mV |
| $\tau_w$ | 120 ms |
| $a$ | 2 nS |
| $b$ | 0.1 nA |
| $V_r$ | -46 mV |
| $V_s$ | 20 mV |

